# *Botrytis cinerea* strains infecting grapevine and tomato display contrasted repertoires of accessory chromosomes, transposons and small RNAs

**DOI:** 10.1101/2022.03.07.483234

**Authors:** Adeline Simon, Alex Mercier, Pierre Gladieux, Benoît Poinssot, Anne-Sophie Walker, Muriel Viaud

## Abstract

The fungus *Botrytis cinerea* is a polyphagous pathogen that encompasses multiple host-specialized lineages. While several secreted proteins, secondary metabolites and retrotransposons-derived small RNAs have been characterized as virulence factors, their roles in host specialization remain unknown. The aim of this study was to identify the genomic correlates of host-specialization in populations of *B. cinerea* associated with grapevine and tomato. Using PacBio sequencing, we produced complete assemblies of the genomes of strains Sl3 and Vv3 that represent the French populations T and G1 of *B. cinerea*, specialized on tomato and grapevine, respectively. Both assemblies revealed 16 core chromosomes that were highly syntenic with chromosomes of the reference strain B05.10. The main sources of variation in gene content were the subtelomeric regions and the accessory chromosomes, especially the chromosome BCIN19 of Vv3 that was absent in Sl3 and B05.10. The repertoires and density of transposable elements were clearly different between the genomes of Sl3 and Vv3 with a larger number of subfamilies (26) and a greater genome coverage in Vv3 (7.7%) than in Sl3 (14 subfamilies, 4.5% coverage). An Helitron-like element was found in almost all subtelomeric regions of the Vv3 genome, in particular in the flanking regions of a highly duplicated gene encoding a Telomere-Linked Helicase, while both features were absent from the Sl3 and B05.10 genomes. Different retrotransposons in the Sl3 and the Vv3 strains resulted in the synthesis of distinct sets of small RNAs. Finally, extending the study to additional strains indicated that the accessory chromosome BCIN19 and the small RNAs producing retrotransposons Copia_4 and Gypsy_7 are common features of the G1 population that are scarcely if ever found in strains isolated from other populations. This research reveals that accessory chromosomes, repertoires of transposons and their derived small RNAs differ between populations of *B. cinerea* specialized on different hosts. The genomic data characterized in our study pave the way for further studies aiming at investigating the molecular mechanisms underpinning host specialization in a polyphagous pathogen.

## Introduction

While specialist phytopathogenic fungi are highly specific to a single host plant, some other fungi stand out for having a broad host range and are qualified as generalist. Nevertheless, these so-called generalist pathogens could actually correspond to multiple co-existing populations that show a certain level of host specialization (Gladieux et al., 2018; Stukenbrock & McDonald, 2008). Such scenario provides an excellent opportunity to investigate the molecular determinants of host specialization. In this regard, the grey mould fungus Botrytis cinerea has recently become a model of choice as both powerful functional genomic tools and population data are available (Choquer et al., 2007; Mbengue et al., 2016; van Kan et al., 2017; Veloso & van Kan, 2018). This ascomycete species is a necrotrophic pathogen that infects more than 1400 host plant species belonging to 580 genera causing significant damages in grapevine and in most cultivated fruits (Elad et al., 2015). Several studies conducted in different parts of the world revealed that populations of B. cinerea are structured, and the host plant was recognized as the factor with the highest explanatory power for this structure, ahead of geography (reviewed in Walker, 2015). Notably French populations of B. cinerea isolated from tomato and grapevine are differentiated from each other (Walker et al., 2015), and have higher aggressiveness on their host-of-origin than other strains, indicating host specialization (Mercier et al., 2019). Recently, Illumina sequencing of 32 representative isolates confirmed the subdivision of these B. cinerea French populations into two genetic clusters on grapevine (G1 and G2 populations) and another, single cluster on tomato that diverged from the G2 population (T population; Sup. Fig. S1; Mercier et al., 2021). These genomic data also allowed to investigate the molecular differences underlying host specialization. By characterizing single-nucleotide polymorphisms in Illumina short-read data, genes with footprints of positive selection and/or divergent selection in the genomes of populations specialized to different hosts were identified. These candidate genes that represent possible determinants of host specialization were enriched in genes encoding Plant Cell Wall Degrading Enzymes (PCWDEs) and transporters and in genes involved in the oxidative stress response. The same study also highlighted a limited number of candidate genes that are population-specific including a couple of PCWDE-coding genes and one secondary metabolism gene cluster (Mercier et al., 2021). Though this previous study provided significant information about the evolution of B. cinerea genes in populations and identified candidate genes possibly involved in host specialization, the lack of a complete assembly of the sequenced genomes did not allow to investigate other mechanisms potentially involved in specialization, such as (i) variation in the presence of accessory chromosomes, (ii) chromosomal rearrangements, and (iii) differences in repertoires of transposable elements (TEs). This last point is of particular interest since retrotransposons were shown to be the source of synthesis of small interfering RNAs (siRNA) acting as effectors in B. cinerea. Indeed, these fungal siRNAs highjack the silencing machinery of the plant cell to reduce the expression of genes involved in the defence process (Weiberg et al., 2013; Porquier et al., 2021). High variability in the repertoire of TEs across fungal populations and the fact that retrotransposons are the main source of siRNAs raise the possibility that these TEs could be involved in host specialization.

In this study, our objective was to investigate whether strains from French populations of B. cinerea specialized on tomato versus grapevine differ in genomic content in terms of core and accessory chromosomes, TEs and small RNA repertoires. We characterized genome sequences produced with the PacBio technology for two representative strains: the Sl3 strain that belongs to the T population specialized on tomato and the Vv3 strain that belongs to the G1 population specialized on grapevine. The full assembly of Sl3 and Vv3 genomes provided a complete view of the core and accessory chromosomes as well as the full repertoires of genes and TEs. Small RNAs produced by the repertoires of retrotransposons were also compared. Finally, extending the study to additional strains allowed to identify which of the genomic features are associated to the populations specialized on tomato or grapevine.

## Results

### The genome architecture of the *Botrytis cinerea* strains Sl3 and Vv3 differ from the B05.10 strain in their accessory chromosomes

Genome assemblies based on PacBio Sequel sequencing data identified 16 core chromosomes (CCs) and two accessory chromosomes (ACs) in both *Botrytis cinerea* strains Sl3 and Vv3 collected on tomato and grapevine. Sequencing reads were assembled *de novo* using a combination of HGAP4 (SMRT-Link v5.0.1) (https://github.com/PacificBiosciences/) and Canu v1.6 (Koren et al., 2017). Both Sl3 and Vv3 assemblies consisted of 18 main contigs. The genome assemblies were estimated to be 43.2 Mb (Sl3) and 44.9 Mb (Vv3), which is close to the 42.6Mb of the B05.10 genome (van Kan et al., 2017). Contigs of the new genome assemblies were ordered and oriented as the previously described chromosomes of B05.10 (BCIN01 to BCIN18) (Fig. 1), and then compared to the genome of B05.10 with Quast (Gurevich et al., 2013) (Sup. Table S1). The length (1.9-4.2 Mb) and GC content (41-43%) of the 16 largest contigs of Sl3 and Vv3 were similar to those of the 16 CCs defined in B05.10. In addition, the high percentage of B05.10 genome coverage (94-99%) confirmed that the 16 largest newly sequenced contigs corresponded to CCs BCIN01 to BCIN16 of *B. cinerea*. The length of CCs was homogenous across the three strains, with less than 400 kb variation per CC (BCIN03: 3.6 Mb in Vv3, 3.3 Mb in Sl3 and 3.2 Mb in B05.10). The length (0.2-0.5 Mb) and GC content (<40%) of the two smallest contigs of Sl3 and Vv3 was typical of ACs, as those found in B05.10. The percentage of B05.10 genome coverage indicated that both strains had the AC BCIN17 (coverage >96%) while only Sl3 had an AC related to BCIN18 (52% coverage). The second putative AC of Vv3 did not share any similarity with the genome of B05.10 and corresponded to a new AC that we numbered BCIN19. Separation of chromosomes by pulse-field gel electrophoresis confirmed the presence of two ACs in each strain (Sup. Fig. S2). According to the migration, the two smallest chromosomes of the Sl3 strain had sizes that corresponded to the assembled contigs BCIN17 and BCIN18 (299 and 188 kb). The same congruence was also observed for the AC BCIN17 of the Vv3 strain (315 kb). In opposite, the migration of the second small chromosome of Vv3, BCIN19, suggested that approximately 100 kb were missing in the corresponding contig (534 kb).

**Figure 1:**
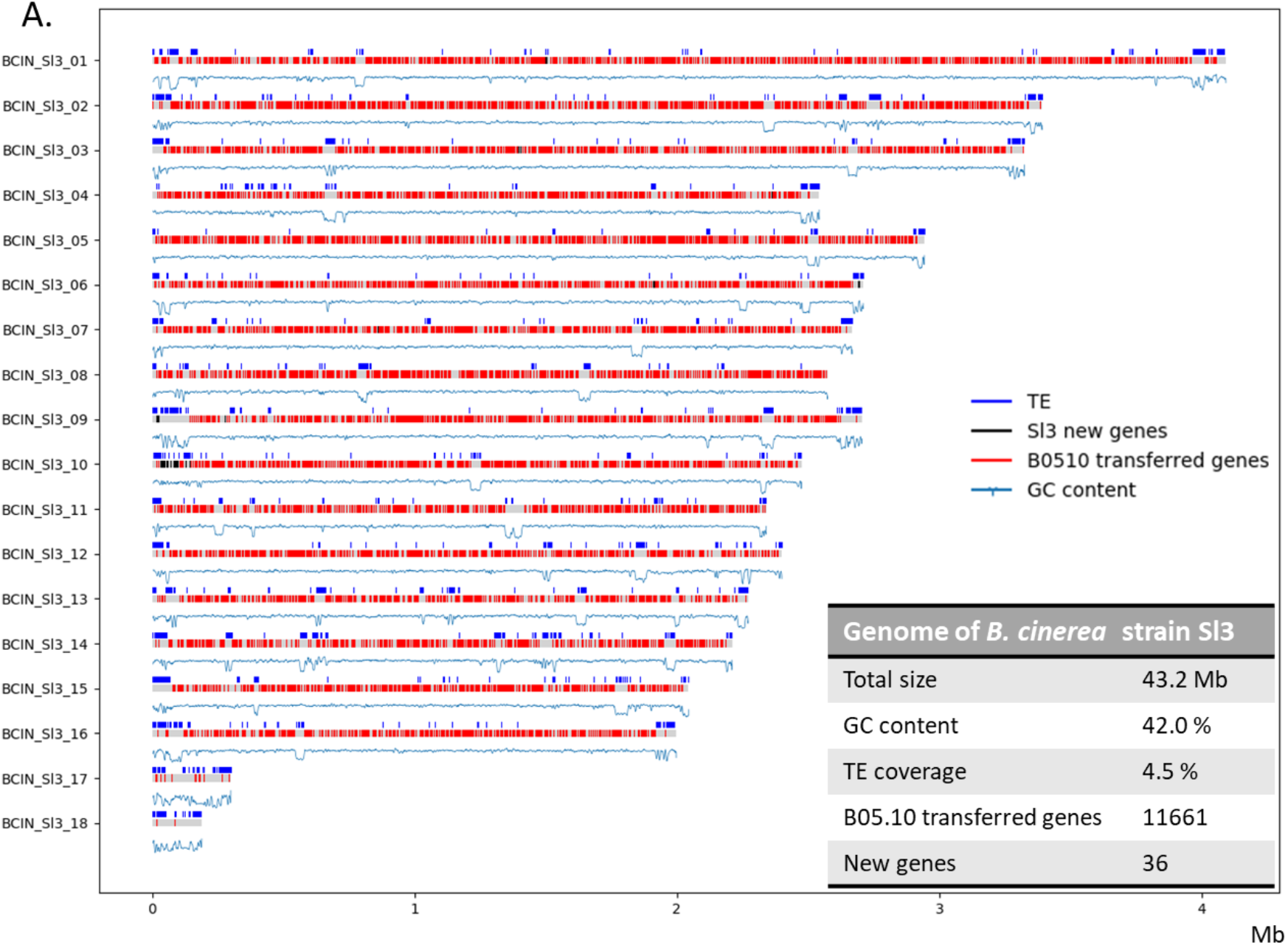
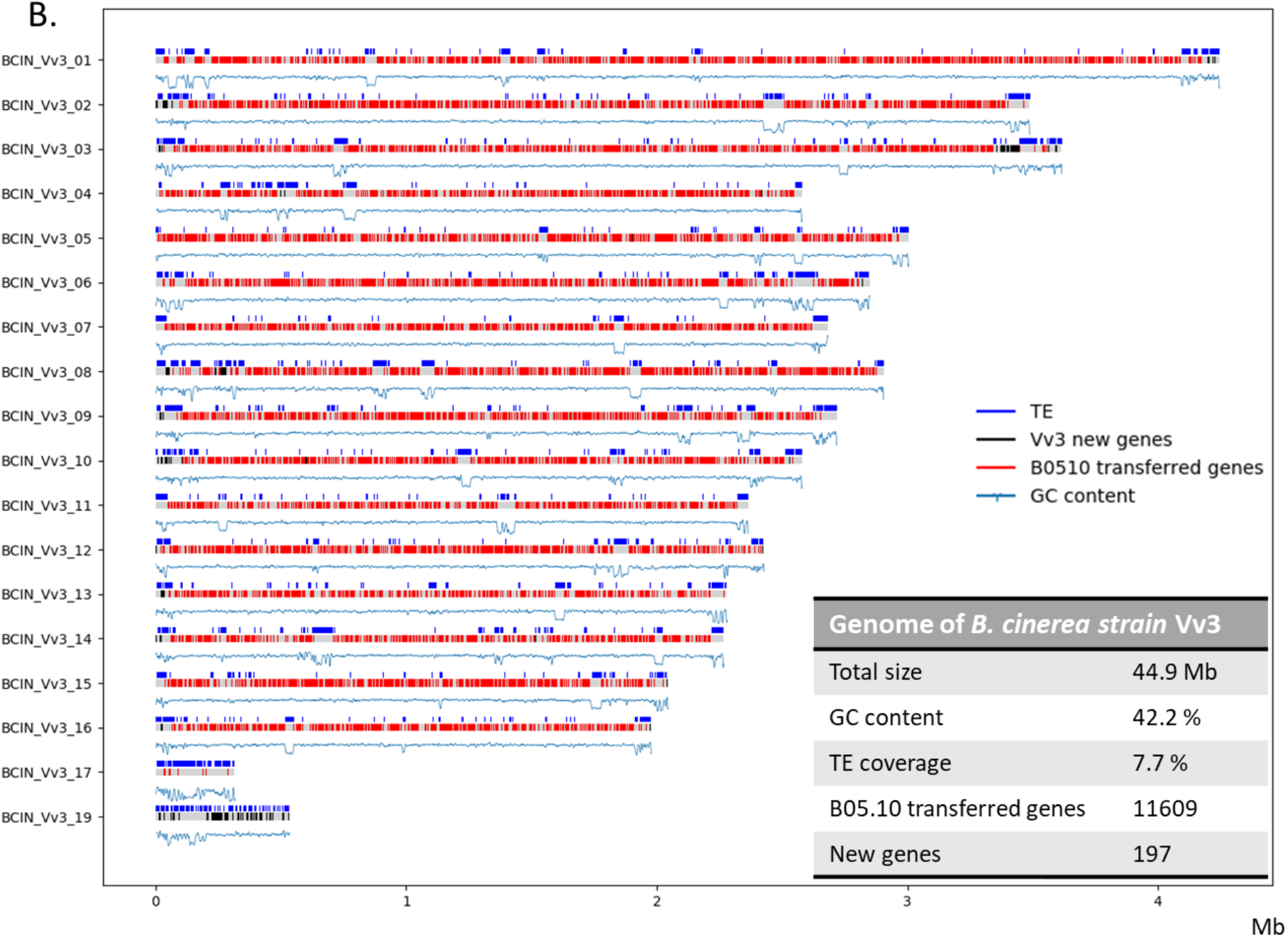
Genome organization in *B. cinerea* strains Sl3 (A) and Vv3 (B). The two karyoplots show chromosomes identified in each strain, genes previously predicted in the B05.10 reference strain (van Kan et al., 2017; red), newly detected genes (black), Transposable Elements (TEs; dark blue) and the GC content (blue) along the Core Chromosomes (CCs BCIN_01 to 16) and the Accessory Chromosomes (ACs BCIN_17 to 19).

A criterion to evaluate the quality of a genome assembly is the presence of telomeres at the terminal regions of the putative full chromosomes. For Sl3, telomeric repeats (*i*.*e*. TTAGGG repetitions; Levis et al., 1997a) were found at both ends of eight chromosomes and at one end of eight other chromosomes (Sup. Table S1). Only the assemblies of CCs BCIN1 and BCIN12 lacked both telomeric repeats. Thus, telomeric repeats were missing at twelve chromosomal ends in the Sl3 assembly. In comparison, telomeric repeats were missing at nine chromosomal ends in the B05.10 assembly (van Kan et al., 2017), though not necessarily at the same chromosomal ends as in Sl3. Telomeric repeats in Vv3 contigs were retrieved only at the 5’ end of CC BCIN2. To investigate the relative scarcity of telomeric repeats in the Vv3 assembly, we searched the raw reads for telomeric repeats. We found 317 and 3712 occurrences of the telomeric repeat (TTAGGG)_3_ and its reverse complement in Vv3 and Sl3 reads, respectively. The relatively low abundance of telomeric repeats in sequencing reads of Vv3 indicated that telomeric sequences were relatively rare in the library. This could be due to degradation of these sequences during the preparation of libraries, as previously described (Tolios et al., 2015).

As the assemblies of the Vv3 and Sl3 genomes revealed different pairs of ACs, we further investigated whether there was a correlation between the content in ACs and the host of origin. We used the 35 available genomic sequences of *B. cinerea* strains from three distinct populations specialized on tomato (T population) or grape (G1 and G2 populations) as well as those of additional strains isolated from bramble (Rf1, Rf2), hydrangea (Hm1), tomato (T4) and grapevine (BcDW1) (Mercier et al., 2021 ; Amselem et al., 2011; Blanco-Ulate et al., 2013). Percentage of chromosome coverage showed different distributions of the three ACs BCIN17, BCIN18 and BCIN19 (AC17 to 19; Table 1). The AC BCIN17 first identified in B05.10 was detected in 30 other strains, including strains belonging to populations T, G1 and G2 as well as in Rf1, Hm1 and BcDW1. The AC BCIN18 also identified in B05.10 was detected in seven other strains, including strains belonging to the T and G2 populations, Hm1 and BcDW1. The AC BCIN19 identified in the Vv3 strain, was specific to the G1 group (detected in 11 strains out of 12). Among populations T, G1 and G2, only G1 showed a relatively homogenous content in ACs with the systematic presence of BCIN17, very high frequency of BCIN19 and absence of BCIN18.

**Table 1:**
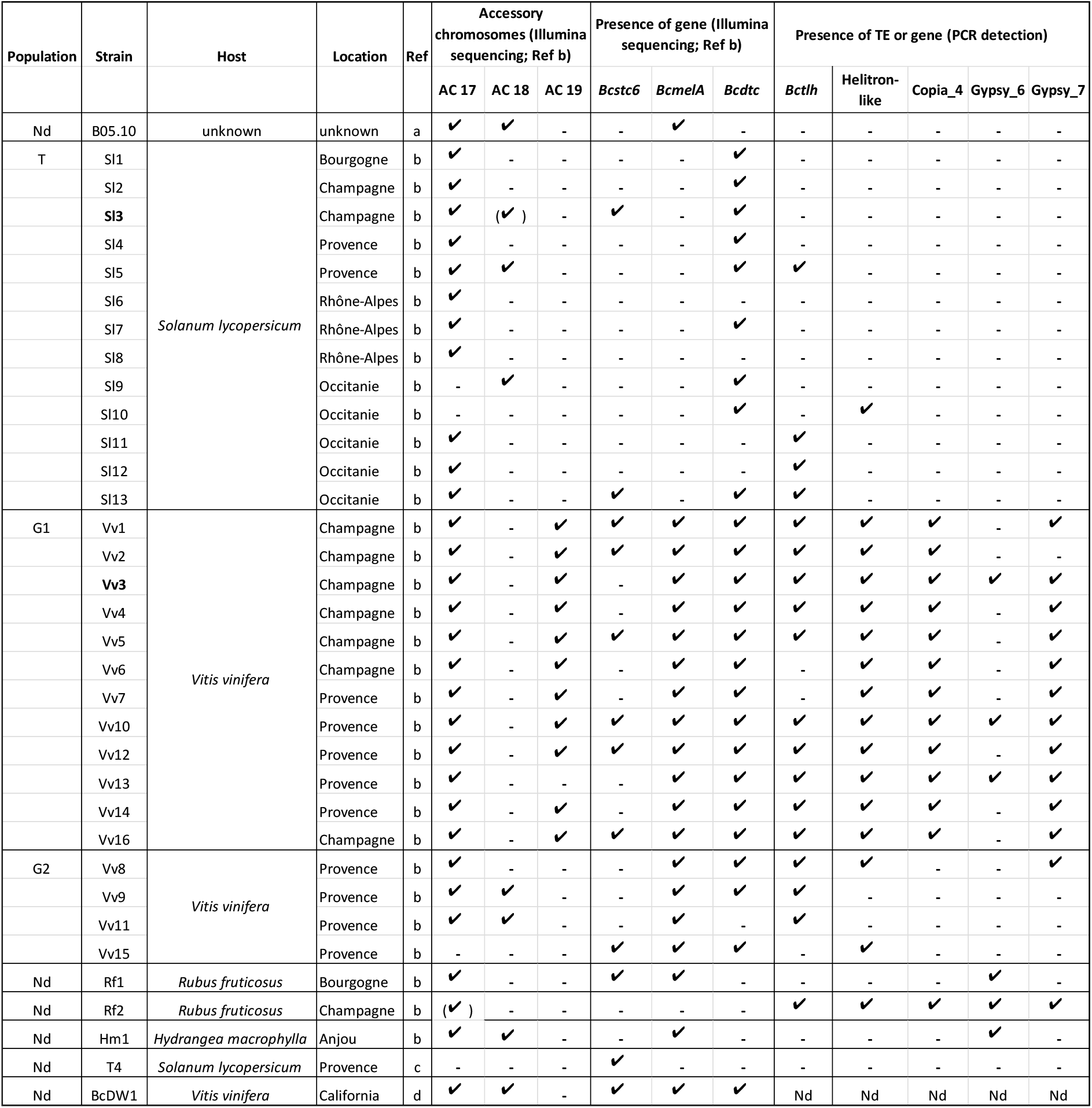
Distribution of Accessory Chromosomes (ACs) and a selection of dispensable genes and Transposable Elements (TEs) in *B. cinerea* strains from different populations. The genomic features highlighted in the present study, using complete assemblies of Sl3 and Vv3 genomes, were investigated in populations specialized either on grapevine (G1 and G2) either on tomato (T; Mercier et al., 2021). The distribution of ACs and secondary metabolism key genes were investigated using Illumina genomic data (Mercier et al., 2021) while those of the TEs and of the gene encoding the telomere-linked helicase (BcHTL) were investigated by PCR. a. van Kan et al., 2017. b. Mercier et al., 2021. c. Amselem et al., 2011. d. Blanco-Ulate et al., 2013. Nd, not determined. ✓, presence. -, absence. (✓), partial presence of an AC. Nd, Not determined.

### The core chromosomes of B05.10, Sl3 and Vv3 strains show a high level of synteny

A vast majority of the genes identified in the reference strain B05.10 were retrieved in the Sl3 and Vv3 genomes. B05.10 gene annotation (van Kan et al., 2017) was transferred to Sl3 and Vv3 with the Liftoff annotation mapping tool (Shumate & Salzberg, 2021). 11 661 and 11 609 genes were transferred to Sl3 and Vv3 respectively, *i*.*e*. 99.6% and 99.2% of the 11 707 genes previously predicted in B05.10. Altogether, 11 590 genes were retrieved in all three genomes, the vast majority in CCs (11 572 genes) and very few in ACs (BCIN17: 18 genes). To assess synteny, we used SynChro (Drillon et al., 2014), a tool that reconstructs synteny blocks between pairwise comparisons of multiple genomes. As shown in Sup. Fig. S3, the three sets of CCs were largely syntenic. When considering major synteny breaks, *i*.*e*. those involving at least four adjacent genes that were not organized as in the B05.10 reference genome, only one and two possible inversion events could be identified in Sl3 and Vv3, respectively (Sup. Fig. S4). Two rearrangements between the terminal regions of chromosomes were also observed, both accompanied by gene duplication. The first one involved a secondary metabolism gene cluster with the polyketide synthase BcPKS7 as a key enzyme (Bcin10g00010-40). This cluster and the two 3’ flanking genes were located in two different regions in B05.10 and Vv3, *i*.*e*. the 5’ end of BCIN10 in B05.10 *versus* the 3’ end of BCIN03 in Vv3. (Sup. Fig. S5). The rearrangement also resulted in the duplication of a gene encoding a putative mitochondrial NADPH cytochrome b5 reductase in Vv3. The second major subtelomeric rearrangement was observed when comparing the genomes of B05.10 and Sl3. The nine first genes of BCIN08 in B05.10 (Bcin08g00010-90) were localized at the start of chromosomes 2 and 15 in Sl3, while BCIN08 in Sl3 started with Bcin08g00080 (Sup. Fig. S6). Hence, the genome of Sl3 contained two copies of seven of these genes and three copies of two of them. The duplicated genes were predicted to encode a P450 monooxygenase, a laccase (*Bclcc11*), a glycoside hydrolase possibly acting on hemicellulose or pectin chains (GH43 family; Lombard et al., 2014), a glycosyl transferase, a heterokaryon incompatibility protein, a pyridine transferase as well as two secreted proteins with unknown functions.

### Subtelomeric regions of core chromosomes and accessory chromosomes provide variation in gene content

The CCs of *B. cinerea* showed Presence Absence Variation (PAV) of genes in particular for secondary metabolism gene clusters. To identify genes present in the Sl3 or Vv3 genomes but not in the B05.10 genome, we used the Fgenesh *ab initio* gene-finder (Solovyev et al., 2006) and excluded the gene models mapping in the B05.10 genome (Sup. Table S2). As shown in Fig. 1, *de novo* genes annotation identified new genes in the CCs especially in their subtelomeric regions. When considering groups of contiguous genes that were not shared between the three genomes, three secondary metabolism gene clusters were identified based on the presence of genes encoding key enzymes. The first one was only detected in Sl3 and included the gene encoding the previously identified SesquiTerpene Cyclase BcSTC6 (BCIN_Sl3_03_394) (Amselem et al., 2011). The second gene cluster was identified in a subtelomeric region of Vv3 and B05.10 (Bcin02g00013 to Bcin02g00016) and included a gene encoding a non-ribosomal peptide synthetase-like similar to MelA, an enzyme involved in the biosynthesis of an α-keto acid dimer in *Aspergillus terreus* (Geib et al., 2016). Finally, the third gene cluster was in a subtelomeric region shared by Sl3 and Vv3 (BCIN_Sl3_10_11 to 26; BCIN_Vv3_10_2 to 14) and included a gene predicted to encode a DiTerpene Synthase (DTC). When looking into previous Illumina sequencing data (Mercier et al., 2021), these three secondary metabolism genes clusters had different distributions: while the *Bcstc6* and the newly identified *Bcdtc* clusters were detected in strains of the three different populations, the *BcmelA* cluster was present in all strains of the G1 and G2 groups but absent from the T group (Table 1).

Fourteen subtelomeric genes coding for a protein predicted to be a helicase were identified in the genome of Vv3 strain but not in the ones of B05.10 and Sl3 strains. This protein showed 51% similarity with the Telomere-Linked Helicases (TLHs) identified in *Magnaporthe oryzae* (Gao et al., 2002; Rehmeyer et al., 2009) and in *Ustilago maydis* (Sánchez-Alonso & Guzmán, 1998). It also shared the same predicted domains *i*.*e*. two C2H2 zinc finger motifs, a helicase ATP-binding domain, a helicase C domain and a specific TLH domain (Rehmeyer et al., 2009) ; Sup. Fig. S7). As in *M. oryzae* and *U. maydis*, the 14 Vv3 helicase-encoding genes were localized in subtelomeric regions (11 regions). They were located between 9 bp and 63 Kbp from chromosome ends, and six of them were actually the first genes detected at the end of the chromosome. Incomplete copies of these helicase-encoding genes were also detected in eight other subtelomeric regions (Sup. Fig. S8). Because of their homology with TLHs of other fungi and of their genomic localization, these newly identified helicases were called BcTLHs. PCR screening of *B. cinerea* populations further indicated that one or several copies of *Bctlh* are present in a majority of strains specialized on grapevine, but also in few strains specialized on tomato or isolated from other hosts (Table 1).

Accessory chromosomes were an additional source of PAV of genes among the B05.10, Sl3 and Vv3 strains (Fig. 1; Sup. Table S1). Firstly, the AC BCIN18 that was present in both Sl3 and B05.10 strains was found to be only partially conserved as it carried seven genes that were shared between the two strains, and nine contiguous genes that were present only in B05.10. Secondly, the newly identified AC BCIN19 of the strain Vv3 carried 78 genes that were not present in the Sl3 and B05.10 strains. These 78 genes mostly encoded proteins with unknown functions, which is reminiscent of the gene content of ACs BCIN17 and BCIN18 (van Kan et al., 2017). Nevertheless, a high proportion of the predicted proteins displayed InterPro (IPR) domains that could be related to nucleic acids binding and modifications (eight ribonucleases, two helicases, two zinc finger C2H2 proteins), transport through vesicles (five dynamins) and peptidase activity (five peptidases) (Sup. Table S2).

### B05.10, Sl3 and Vv3 strains show different repertoires and densities of transposable elements

Different subfamilies of Transposable Elements (TEs) were identified in the B05.10, Sl3 and Vv3 strains. The complete annotation of TEs in the Sl3 and Vv3 genomes was carried out using a *de novo* approach with the REPET package (Flutre et al., 2011; Amselem et al., 2015). Consensus sequences representing all possible TEs among the Sl3 and Vv3 genomes were identified. After manual curation, 19 and 33 consensus TE sequences were retained for Sl3 and Vv3, respectively (Sup. File S1). They were classified based on their structure and sequence similarities to characterized eukaryotic transposons (Hoede et al., 2014; Wicker et al., 2007). Fifteen consensus sequences were previously identified in the B05.10 genome using the same pipeline and parameters as in the present study (Porquier et al., 2016). Comparison of the B05.10, Sl3 and Vv3 consensus TE sequences revealed a total of 33 different subfamilies: 13 class I LTR (retrotransposons), seven class II TIR (DNA transposons), one class II Helitron and 12 subfamilies with either low genome coverage or vague classification (Fig. 2; Sup. Table S3). The subfamily *Boty*/Gypsy_1 element (Diolez et al., 1995) was built from several *Boty* consensus as they were identical in their central part and only differed at terminal regions (Sup. Fig. S9), as previously observed in B05.10 (Porquier et al., 2016, 2021). In total, eight subfamilies were shared between the three genomes, *i*.*e*. two subfamilies of Copia retrotransposons, four subfamilies of Gypsy retrotransposons including the *Boty*/Gypsy_1 element and two subfamilies of the Tc1-Mariner DNA transposons, including the *Flipper* element (Levis et al., 1997b). Fifteen, four, and three subfamilies were exclusive to the Vv3, Sl3 or B05.10 genomes, respectively, and three subfamilies were shared between two genomes. Libraries of subfamily sequences were used to annotate TE copies in the whole genomes. For each subfamily, both full-length and truncated copies were retrieved in the genomes.

**Figure 2:**
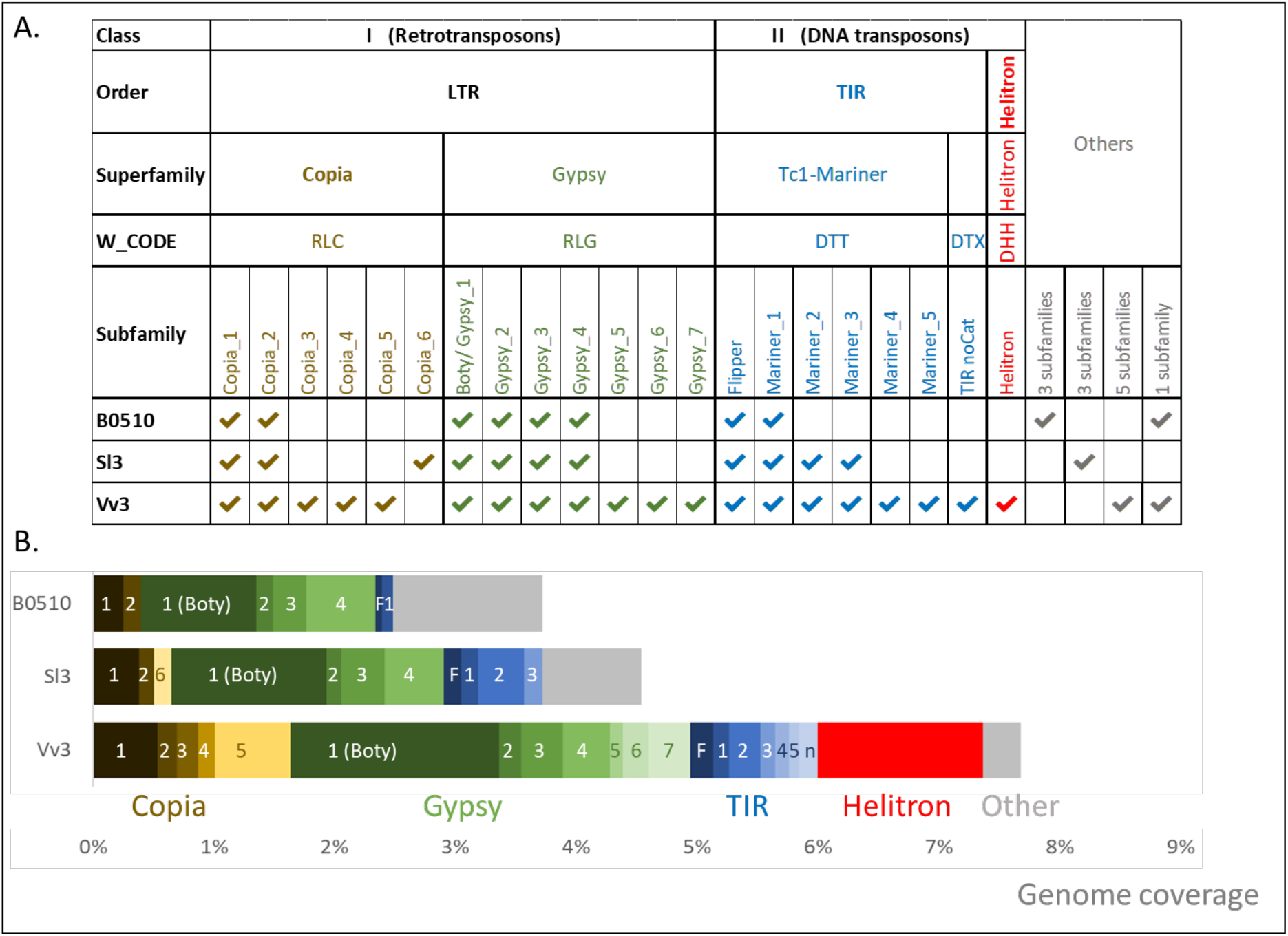
Transposable Elements (TEs) in the genomes of the *B. cinerea* strains B05.10, Sl3 and Vv3. A. Subfamilies of TEs identified in the Sl3 and Vv3 strains were classified according to Wicker et al., (2007), as previously done for the strain B05.10 (Porquier et al., 2016). In the present study, all *Boty* consensus were merged into a single subfamily. B. Total genome coverage was detailed for the different subfamilies. For more details, see Sup. Table 3.

Genome coverage by subfamilies of TEs differed among the strains. Percentages of coverage for each subfamily in each genome were computed (Fig. 2; Sup. Table S3). Total coverage by TEs was higher for the Vv3 genome (7.7%), in which the largest number of subfamilies was identified, than in Sl3 and B05.10 (4.5% and 3.7%, respectively) (Porquier et al., 2016). Transposable Elements were detected at all chromosome ends but the 3’ end of BCIN08 in Sl3 (Fig. 1), and also inside chromosomes, with a significantly higher coverage in ACs (27.9-35.4%) than in CCs (1.9-11.3%). The eight shared subfamilies were responsible for about 3% of the coverage in each genome whereas other subfamilies covered approximately 1% of B05.10 or Sl3 genomes and 3.6% of Vv3 genome. The specific subfamilies responsible for the higher coverage in Vv3 were mainly Copia and Gypsy retrotransposons and a TE that was similar to a Helitron and covered 1.37% of the Vv3 genome.

The subtelomeric regions of the chromosomes of the Vv3 strain were invaded by a Helitron-like TE. Helitron is a unique class of DNA transposons in eukaryotes and the representative elements of this class encode a protein with two enzymatic domains corresponding to the helicase and the nuclease activities that are required for transposition (Kojima, 2019). As only the helicase domain was clearly identified in the TE found in Vv3, it was called a Helitron-like element (Sup. Fig. S10). Helitrons are also known for their capacity to capture genes, which increases their size (Castanera et al., 2014). In addition to a helicase-coding gene, the 15.5 kb sequence of the Helitron-like element found in Vv3 revealed two captured genes as well as relics of *Boty*/Gypsy_1 (*i*.*e*. Long Terminal Repeats). For one of the captured genes, the predicted protein was a secreted pectate-lyase (IPR002022), similar to enzymes involved in the maceration and soft rotting of plant tissue. Notably, 33 complete or uncomplete copies of the Helitron-like TE carried this putative virulence gene in the genome of Vv3. Unlike most TEs that were randomly distributed along chromosomes, the Helitron-like TEs were mainly localized at the chromosome ends (Sup. Fig. S11). Looking closer at their precise localization, we noticed that they were found in 3’ and/or 5’ of all the 14 complete copies of the helicase encoding gene *Bctlh* (Sup. Fig. S8). Finally, to investigate how this Helitron-like TE was distributed among the populations of *B. cinerea*, we designed PCR specific primers. The results indicated that all G1 strains specialized on grapevine carried one or several copies of the Helitron-like TE while the element was scarcely detected in the T population and detected in two strains out of four in the G2 population (Table 1).

Fungi have developed defense mechanisms against invasion by TEs, including a Repeat-Induced Point mutation (RIP) machinery that inactivates repeated sequences by causing Cytosine to Thymine mutations and therefore decreases GC content. In a previous study, Amselem et al. (2015) showed that the genome of *B. cinerea* contains the genes encoding the cytosine methyltransferases that are required for RIP (BcRID1 and BcRID2), as well as TEs with signatures of RIP at both CpT (10-40% of TEs) and CpA loci (10-35% of TEs). However, depending on the TE copy and on the *B. cinerea* strain, the occurrence of RIP seems to be highly variable (Porquier et al., 2016, 2019, 2021). Regarding the complete copies of TE from Sl3 and Vv3 (Fig. 3; Sup. Fig. S12), the majority of the subfamilies displayed copies with a large range of GC contents. For example, the Flipper DNA transposon showed both copies with a GC content similar to those previously observed in a mobile copy (39%; Levis et al., 1997b) and copies with a lower GC content (<20%) which could suggest inactivation by RIP. For each subfamily of class I and II TEs identified in B05.10, copies with different GC content were observed. The same pattern was observed for Sl3 except for subfamily Copia_2 for which all four copies had a GC content > 40%. In contrast, the Vv3 genome included seven subfamilies of TEs, *i*.*e*. the Copia_2 and 4, Gypsy_5, 6 and 7, Mariner_4 and Helitron-like subfamilies, that contain only copies with GC content >40% indicating the absence of RIP. Apart from Copia_2, these subfamilies are absent in the Sl3 and B05.10 genomes.

**Figure 3:**
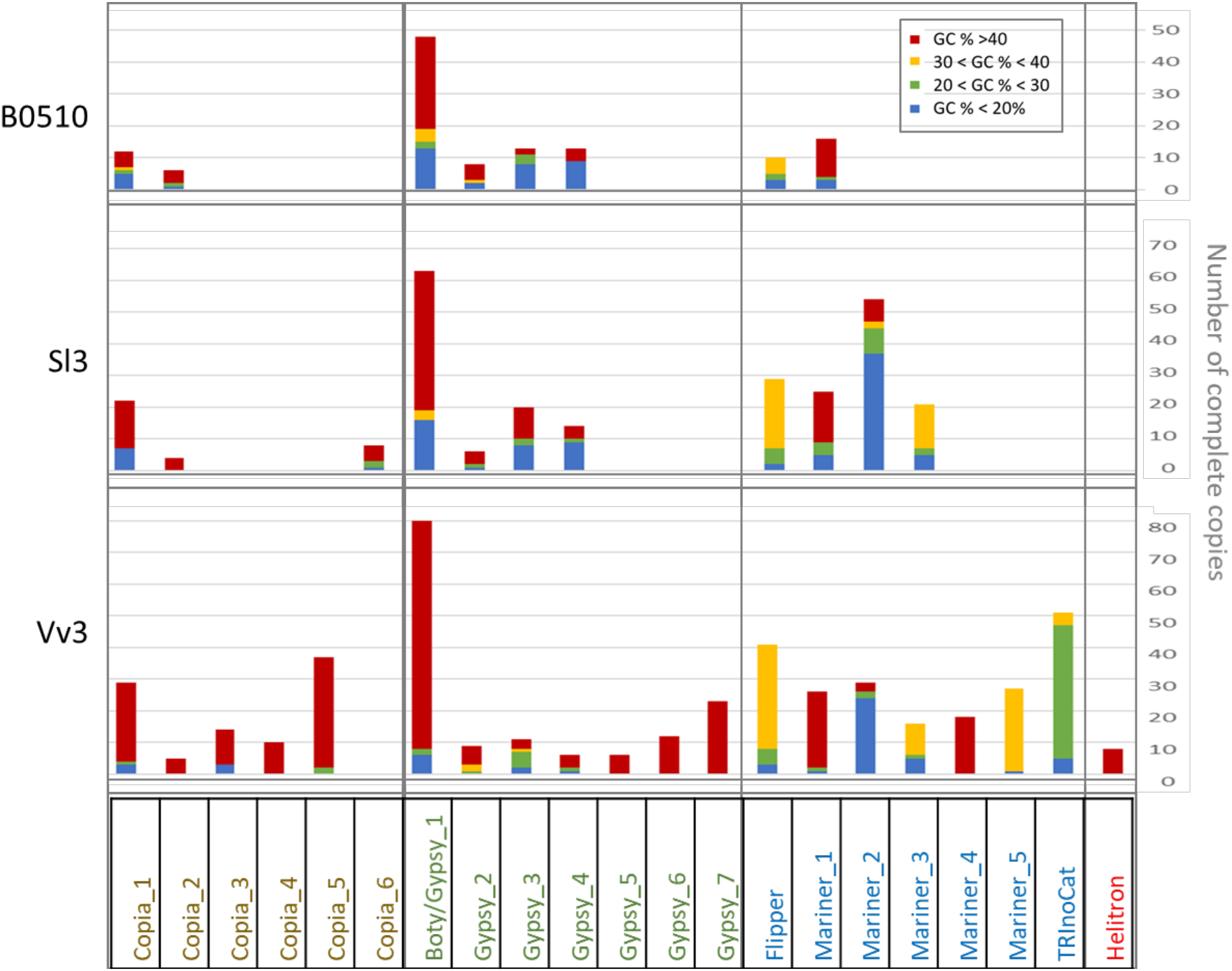
GC content of complete copies of the main Transposable Elements (TEs) in the genomes of *B. cinerea* strains B05.10, Sl3 and Vv3. The complete copies of each subfamily of TEs were classified according to their GC content. See Sup. Fig. S11 for more details.

### Retrotransposons of the Sl3 and Vv3 strains produce different sets of small RNAs

As retrotransposons and especially unripped copies are known to be the origin of the synthesis of small RNAs in *B. cinerea* (Weiberg et al., 2013; Porquier et al., 2021), we investigated whether the different repertoires of TEs in the Sl3 and Vv3 strains could lead to the production of different sets of small RNAs. The two strains were grown both on grape and tomato juice solid media to partially mimic the conditions that the fungus encounters on the host plants (Simon et al., 2013). The corresponding small RNA libraries were Illumina-sequenced. After quality filtering (Sup. Table S4), only the reads between 20 and 24 nucleotides were kept as they correspond to the size of the small silencing RNAs generated by the cleavage of long double stranded RNAs by the Dicer nuclease (Weiberg et al., 2013). A Principal Component Analysis (PCA) of the repertoires of small RNAs in the four samples indicated that the two strains harbour different repertoires, and that the use of grape *versus* tomato juice culture medium had little impact on those repertoires (Sup. Fig. S13, A).

In order to identify which TEs were involved in the production of small RNAs, the reads from the Sl3 and Vv3 strains were mapped against the consensus sequences of the 33 identified TE subfamilies. The results were similar with the two culture media and indicated that the TE-derived small RNAs were produced by retrotransposons corresponding to seven subfamilies (Table 2, grey lines):

- Three out of six retrotransposons shared between Sl3 and Vv3 (*Boty*/Gypsy_1, Gypsy_3 and Gypsy_4) produced small RNAs in both strains. These three retrotransposons were previously identified as the only ones producing significant amounts of small RNAs in the B05.10 strain (Porquier et al., 2021).
- The unique retrotransposon found in Sl3 but not in Vv3 (Copia_6) produced small RNAs.
- Three out of six Vv3-specific retrotransposons found in Vv3 but not in Sl3 (Copia_4, Gypsy_6 and Gypsy_7) produced small RNAs.

**Table 2:**
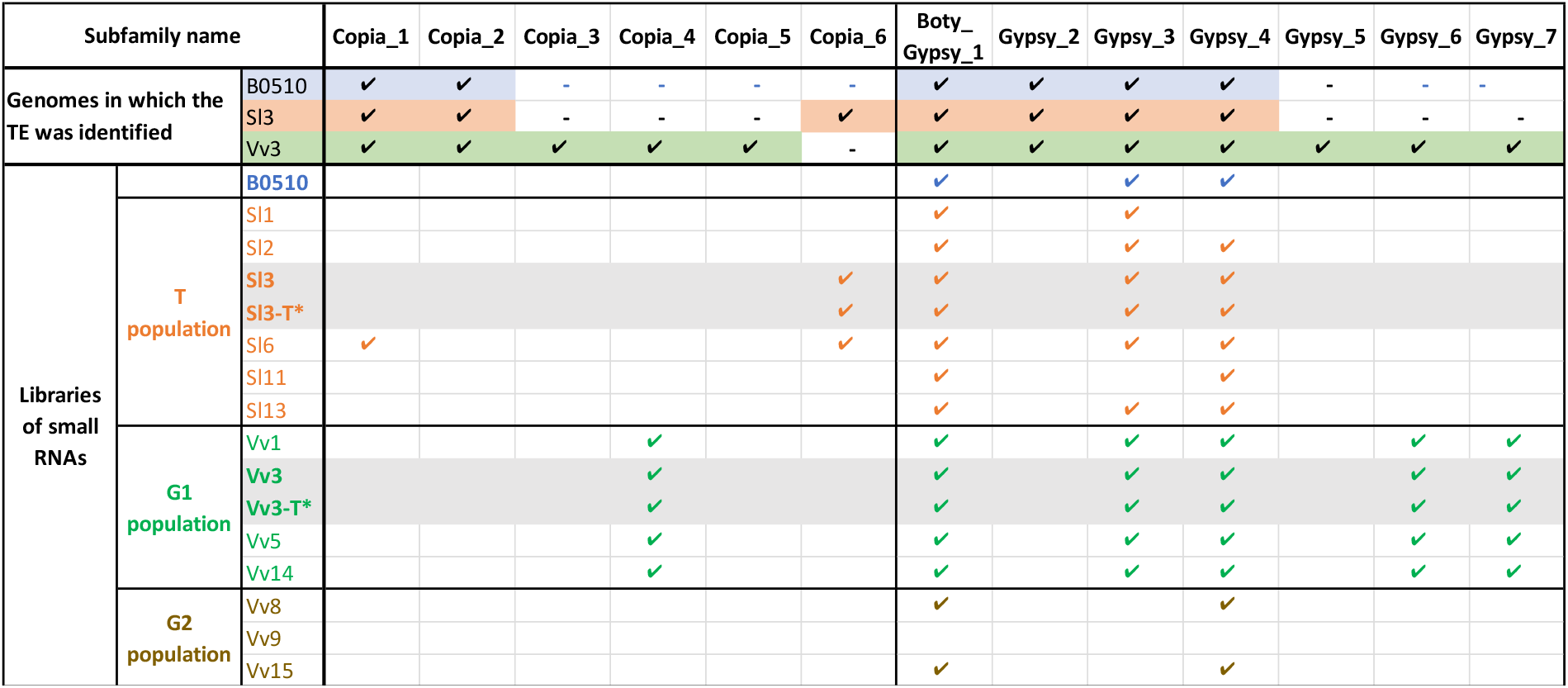
Small RNAs produced by the Transposable Elements (TEs) identified in the *B. cinerea* strains B05.10, Sl3 and Vv3. Small RNA libraries were made from strains cultivated on grape juice medium except two that were done from strains cultivated on tomato juice medium (indicated by -T*). Reads from the 16 libraries were mapped on the TEs to identify the small RNA-producing retrotransposons. For small RNAs, ✓ indicates that at least 300 Reads Per Million (RPMs) from the library are mapping on the corresponding TE. Genomes in which the TEs were identified are reminded at the top of the table (✓, presence. -, absence). For more details, see Sup. Table S4.

In addition to these qualitative differences, quantitative differences were also observed with Gypsy_3 producing the highest amount of small RNAs in the Sl3 strain (291.074 and 283.036 Reads Per Millions (RPMs) in the grape and tomato juice media, respectively (Sup. Table S4) and Copia_4 producing the highest amount of small RNAs in Vv3 strain (115.863 and 168.212 RPMs in the grape and tomato juice media, respectively).

Sequences of the seven small RNA-producing TEs all showed the expected structure for Copia and Gypsy classes of transposons (Wicker et al., 2007), except Gypsy_4 that lacked the GAG domain and Copia_4 that had a small additional one (Sup. Fig S14). Small RNAs mapped almost all along the sequences suggesting that the whole elements could be converted into small RNAs (except from the GAG domain of Gypsy_3). All seven small RNA-producing TEs were in numerous complete copies (six to 80; Sup. Fig. S12) all over the chromosomes of Vv3 or Sl3 strains. The three Vv3 specific small RNA-producing TE, *i*.*e*. Copia_4, Gypsy_6 and 7, only showed copies with relatively high GC content (44.4% +/-0.0, 47.4% +/-0.1 and 43.6% +/- 0.0 respectively). For the other small RNA-producing TEs, *i*.*e* Gypsy_3, Gypsy_4, *Boty*/Gypsy_1 and Copia_6, copies with varied levels of GC were identified. Nevertheless, mapping of the small RNA reads on the genomes showed that only copies with relatively high GC content could generate small RNAs (Sup. Fig. S15).

Among *B. cinerea* small RNAs, the three siRNAs previously identified as effectors (*i*.*e*. siR5, siR3.1 and siR3.2) were 21 bp in size and displayed the nucleotide U in first position (Weiberg et al., 2013). We investigated which proportion of the small RNAs produced by the Vv3 and Sl3 strains shared the same features (Sup. Fig. S16). For all seven TEs, the generated small RNAs were mainly composed of 22 nucleotide sequences (54-69%), followed by 21 nucleotide sequences (18-36%). Notably, a vast majority of them (97%) showed an uracil (U) in first position. Finally, the three siRNA previously identified as effectors in the strain B05.10 (*i*.*e*. siR5, siR3.1 and siR3.2) were identified among the four libraries. Mapping data indicated that siR5 was produced by *Boty*/Gypsy_1 while siR3.1 and siR3.2 were produced by Gypsy_3, as previously demonstrated in the B05.10 strain (Weiberg et al., 2013; Porquier et al., 2021).

### A set of retrotransposons-derived small RNAs is associated with the G1 population specialized on grapevine

As the repertoires of TE-derived small RNAs produced by the strains Vv3 and Sl3 were partially different, we further investigated whether there was a correlation between the previously identified populations of *B. cinerea* specialized on grapevine or on tomato (Mercier et al., 2021) and the production of some of these small RNAs. The study was extended to a total of six strains from the T population, four strains from the G1 population and three strains from the G2 population, as well as to reference B05.10. All strains were grown on grape juice medium and small RNAs were investigated as described above. Principal component analysis of small RNA repertoires (Sup. Fig. S13, B) differentiated samples from the G1 population from samples from the G2 and T populations. To compare the repertoires of TE-derived small RNAs from the 14 strains, we mapped the reads on the TEs identified in B05.10, Sl3 and /or Vv3 strains (Table 2; Sup. Table S4). Mapping data indicated that the same seven retrotransposons that produced small RNAs in B05.10, Sl3 or Vv3 strains could be responsible for the production of small RNAs in the other tested strains. An exception was observed for the Vv9 strain as its small RNAs did not map to any of the known TEs. It should be noted here that for Vv9 and the other ten strains where only unassembled genome sequences are available (Illumina data), additional unknown retrotransposons may be involved in the production of small RNAs. From the presented analysis, the following distribution of retrotransposons derived-small RNAs in the different populations could be observed:

- Small RNAs produced by the shared *Boty*/Gypsy_1 and Gypsy_4 TEs were identified in most of the strains from the three populations and in the strain B05.10.
- Small RNAs produced by the shared Gypsy_3 TE were identified in most of the strains of the T and G1 populations and in the strain B05.10, but not in three strains of the G2 population.
- Small RNAs produced by the Sl3-identified Copia_6 TE were only identified in Sl3 and another strain of the T population (Sl6).
- Small RNAs produced by the Vv3-identified Copia_4, Gypsy_6 and Gypsy_7 TEs were exclusively identified in G1 strains. This could explain why the G1 strains separated from the other strains in the PCA analysis (Sup. Fig. S13, B). In the genome of Vv3, Copia_4, Gypsy_6 and Gypsy_7 were detected in 10, 12 and 23 complete copies, respectively (Sup. Fig. S12). Copia_4 was localized on several CCs, while Gypsy_6 and Gypsy-7 were found both on CCs and ACs (Sup. Fig. S17). The G1-specific AC BCIN19 revealed seven complete copies of Gypsy_7.

We further focused on the potential G1-specific small RNAs by investigating the presence of the three corresponding retrotransposons in strains from the T, G1 and G2 populations and strains from other hosts. Copia_4, Gypsy_6 and Gypsy_7 were searched by PCR using one pair of specific PCR primers per TE. These primers were defined in conserved regions of TEs and were expected to amplify at least the unripped copies of TEs containing these regions (Sup. Fig. S18). As shown in Table 1, Copia_4 was detected in all the G1 strains tested (12 strains) but not in the T and G2 populations (13 and 4 tested strains, respectively). Gypsy_7 was detected in all strains of the G1 population but one (Vv2), in one strain of the G2 population (Vv8) but not in the T population. Finally, Gypsy_6 was detected in only three strains of the G1 population (Vv3, Vv10 and Vv13). Unexpectedly, the PCR approach did not allow to detect Gypsy_6 in Vv1, Vv5 and Vv14 strains, whereas small RNAs generated from this TE were isolated from the same strains (Table 2). We therefore investigated the mapping of these small RNAs on the Gypsy_6 TE identified in the Vv3 strain

(Sup. Fig. S18, C). For Vv5, no small RNA read mapped on the loci of the PCR primers allowing for the possibility that these regions were not conserved in Vv5. By contrast, some small RNAs generated by the Vv1 and Vv14 strains mapped to the Gypsy_6 regions where the PCR primers were defined. One remaining hypothesis to explain the absence of PCR product could be the absence of a copy of Gypsy_6 containing both primer regions in Vv1 and Vv4 genomes. In conclusion, while the absence of PCR products should be interpreted with caution, the results indicated that the small RNA generating-TE Copia_4, and to a lesser extent, Gypsy_7, are enriched in the G1 population.

## Discussion

While the genomes of several strains of the polyphagous pathogen *B. cinerea* have already been sequenced allowing studies of genetic variation (Atwell et al., 2015; Mercier et al., 2021), only one gapless genome, whose of the model strain B05.10, was available so far (van Kan et al., 2017). In this study, we used the PacBio technology to sequence the full genomes of the Sl3 and Vv3 strains that represent two populations of *B. cinerea* specialized to tomato (T population) and grapevine (G1 population; Mercier et al., 2021). We also used additional strains of these two populations and strains from a second population specialized to grapevine (G2) to extend our comparative analyses to a larger set of pathogens.

### The full genomic assemblies of *B. cinerea* strains from different populations reveal highly syntenic core chromosomes

The long-reads generated with the PacBio technology allowed us to generate genome assemblies of 43.2 Mb and 44.9 Mb, for the strains Sl3 and Vv3, respectively. These assembly lengths are close to that of the reference strain B05.10 (42.6 Mb). Additionally, almost all genes (>99%) previously annotated in the strain B05.10 were identified in the generated assemblies. As shown by sequence comparison and gene annotation, the three strains share 16 CCs with a high level of synteny with only few events of inversions or translocations. The same number of CCs was also observed in the other genomes of *Botrytis* and Sclerotiniaceae species (Derbyshire et al., 2017; Valero-Jiménez et al., 2020). The high level of synteny between the CCs of *B. cinerea* strains is consistent with their interfertility, as synteny is needed for appropriate chromosome pairing during meiosis. Direct evidence for interfertility comes from a fertile progeny that was obtained by crossing Sl3 and Vv3 (M. Viaud, *unpublished data*). Indirect evidence for interfertility can be found in population genetic analyses, which previously indicated gene flow between the T and G1 groups to which Sl3 and Vv3 belong (Mercier et al., 2021).

### The accessory chromosomes BCIN19 is specific to the *B. cinerea* G1 population specialized on grapevine

The full genomic assemblies generated in this study revealed that Sl3 and Vv3 strains have different pairs of ACs characterized by their small sizes (0.2-0.6 Mbp), their low GC content (25-38%) and a high density of TEs (28-35%). While the Sl3 strain carries two ACs similar to those of B05.10 (BCIN17 and BCIN18), the Vv3 strain carries BCIN17 and a newly characterized AC named BCIN19. Extending our study to 35 *B. cinerea* strains confirmed the dispensability of BCIN17, BCIN18 and BCIN19 and did not support any lineage specificity for BCIN17 and BCIN18. In opposite, our data suggest that BCIN19 is specific to the G1 group specialized on grapevine. Exploring further the distribution and conservation of the ACs in larger samples would allow to define more precisely to which extent some parts of the accessory genome are associated with a specific population. Though the function of fungal ACs remains largely elusive, several studies on phytopathogenic species reported that ACs could carry genes encoding effectors or genes involved in the synthesis of toxic metabolites (Bertazzoni et al., 2018; Meena et al., 2017; Yang et al., 2020). Our study indicates that ACs are an important source of variation in gene content in *B. cinerea*, but at this stage, none of the 78 genes carried by the ACs has a predicted function indicating a direct role in pathogenicity. Accessory chromosomes could also be considered as reservoirs of TEs. Interestingly, BCIN19 carries several copies of the Gypsy_7 retrotransposon that is specific of the G1 population and generates small RNAs.

### The genome of the *B. cinerea* strain Vv3 has a larger repertoire of transposable elements than the Sl3 and B05.10 strains

Transposons are important features of fungal genomes playing key roles in genome structure, genome plasticity (Fouché et al., 2021), and production of small RNAs (Weiberg et al., 2013). TEs identified in the three fully sequenced and assembled genomes of *B. cinerea* strains B05.10, Sl3 and Vv3 cover 3.7, 4.5 and 7.7% of their genomic sequences, respectively. Variation in TE content explains most of the variation in total assembly length, with the genomic sequences varying between 41.0 and 41.4 Mb when TEs are excluded. As already observed in the B05.10 strain (van Kan et al., 2017), TEs identified in the Sl3 and Vv3 strains are frequent in the subtelomeric regions of the CCs, as well as in ACs where they can cover up to one third of the sequence. While previous studies already indicated variation in the presence of TEs (e.g *Boty* and *Flipper*) in *B. cinerea* strains (Amselem et al., 2011; Porquier et al., 2021), our study is the first that compares TEs between fully assembled genomes. Our data clearly show that the repertoires of TEs are very different between the three sequenced strains and 21 new subfamilies of TEs were recovered, mostly from the Vv3 strain. As in many fungi, TEs of *B. cinerea* could be inactivated by RIP (Amselem et al., 2015; Porquier et al., 2016, 2019, 2021). In our analysis, the GC content of TE copies suggests that RIP is less active in the Vv3 strain, consistent with its larger TE content. This larger content is due to retrotransposons, TIR transposons, as well as to a Helitron-like TE. Several subfamilies of the TEs identified in this strain, including retrotransposons and the Helitron-like TE, do not show any traces of RIP and could possibly be able to transpose in new *loci*.

### Rearrangements at chromosome ends are a source of gene gains and losses in *B. cinerea*

Our synteny and PAV analyses indicate that the few rearrangements occurring in the CCs correspond mainly to exchanges of chromosomes ends, and that most gene gains or losses also happen in these TE-rich regions. In many fungi, subtelomeric regions are enriched in secondary metabolism gene clusters (Kjærbølling et al., 2020). Therefore, rearrangements in these regions could contribute to the intraspecific diversity of the biosynthetic clusters (Olarte et al., 2019). In the genome of *B. cinerea*, six out of the approximately 40 identified secondary metabolism gene clusters are in subtelomeric regions (Amselem et al., 2011; van Kan et al., 2017). The present study provides evidence that the evolution of some of these subtelomeric clusters is subjected to chromosomal ends exchange (PKS7 cluster) as well as to gene gains and losses (MelA-like and DTC clusters). In other studies, partial or total loss of the subtelomeric gene cluster responsible for the biosynthesis of the botcinic acid phytotoxin was observed in some rare strains of *B. cinerea* and in other species of *Botrytis* (Plesken et al., 2021; Valero-Jiménez et al., 2020).

In addition to gene gains and losses, our study revealed that duplication of *B. cinerea* genes occurs in subtelomeric regions. For example, the four colocalized genes (Bcin08g00060 to 90) possibly related to detoxification of reactive oxygen species or plant cell wall degradation and previously identified as duplicated in strains specialized to tomato (T group; Mercier et al., 2021) were localized on two different chromosomes ends of the Sl3 strain.

Subtelomeric regions of all three B05.10, Sl3 and Vv3 strains were found to be enriched in TEs, and TEs were detected in the exchanged chromosomal ends mentioned above. Similar observations made in other fungal genomes led to the hypothesis that TEs may be responsible for repeat-driven recombination and gene rearrangements (Fouché et al., 2021; Lind et al., 2017; Olarte et al., 2019). If these exchanges of chromosomes ends are later followed by a sexual cycle in which chromosome pairing occurs between CCs that have different extremities, it could result in gene gain or loss. Indeed, in this scenario, the progeny would include individuals with two copies of one initial subtelomeric region or none of them. This two-steps mechanism could have an important role in the evolution of the repertoire of secondary metabolites produced by *B. cinerea*.

### A helicase encoding gene and a Helitron-like element shape the subtelomeric regions of the genome of the *B. cinerea* strain Vv3

Analysis of the genome of the *B. cinerea* strain Vv3 revealed that subtelomeric regions show a particular structure with the co-occurrence of a Telomere-Linked Helicase (TLH) encoding gene and a Helitron-like TE that were not described so far in this species. Genes encoding TLHs were previously described in the subtelomeric regions of the fungal pathogens *M. oryzae* and in *U. maydis* (Gao et al., 2002; Rehmeyer et al., 2009; Sánchez-Alonso & Guzmán, 1998). The Vv3 genome contains 14 copies of the *Bctlh* gene, all in subtelomeric regions and, strikingly, all flanked in 3’ and/or 5’ by a copy of a Helitron-like TE. Helitrons were described in other fungi *i*.*e. Pleurotus ostreatus* and *Fusarium oxysporum* (Castanera et al., 2014; Chellapan et al., 2016) but, to our knowledge, this is the first study that shows a strong enrichment of this family of TEs in subtelomeric regions. One hypothesis to explain the localization of the Helitron-like TE of *B. cinerea* could be that it preferentially inserts in the flanking regions of the *Bctlh* gene copies. Nevertheless, as no nuclease domain was detected in the sequence of Helitron-like TE, this may question its ability to transpose (Kojima, 2019). Alternatively, the occurrence of the *Bcthl* gene and Helitron-like TE in the subtelomeric regions could arise from recombination between chromosome ends followed by sexual crosses leading to duplications as discussed above. Indeed, when these mechanisms are repeated in successive generations, they could lead to the homogenization of the subtelomeric regions. Previous studies revealed a drastic reduction of the number of Restriction Fragment Length Polymorphisms (RFLPs) in subtelomeric regions of *B. cinerea* strains isolated from grapevine suggesting a possible homogenization of these regions (Levis et al., 1997a). Based on our analysis, one may wonder whether this homogenization is due to the amplification of the *Bcthl* genes and Helitron-like TEs by exchanges and duplications of chromosomes ends.

Another original feature of the 15.5 kb sequence of the Vv3 Helitron-like TE is that it carries captured genes, as in some other Helitrons elements (Castanera et al., 2014). It would be interesting to evaluate the expression of these captured genes especially the one encoding a putative secreted pectate-lyase and to test whether the numerous copies (33) present in the Vv3 genome confer a higher ability to degrade host tissues. Our analysis to *B. cinerea* populations indicated that the Helitron-like TE is present in all the strains of the G1 group, in about half of the strains of the G2 group and in a minority of strains of the T group which further questions its potential role in the interaction with the host plant.

### Both Gypsy and Copia retrotransposons generates small RNAs in *B. cinerea*

*Botrytis cinerea* produces siRNA that are acting as effectors in the host plant where they can highjack the silencing machinery to impair the expression of genes involved in the defence process (Weiberg et al., 2013). As in other fungal models, these small RNAs are mainly synthesized from retrotransposons. The work just published by Porquier et al., (2021) actually identified *Boty*/Gypsy_1, Gypsy_3 and Gypsy_4 as the only TE producing small RNAs in B05.10, and the authors demonstrated that the Gypsy_3 TE could be considered as a pathogenicity factor by itself as the introduction of Gypsy_3 in a strain lacking this TE resulted in the production of siRNA and an enhanced aggressiveness on tomato and on *Arabidopsis thaliana*. In our study, we demonstrated that the Sl3 and Vv3 strains that have very different repertoires of TEs are consequently able to produce different sets of TE-derived small RNAs. Seven retrotransposons, five Gypsy subfamilies but also two Copia subfamilies, were identified as responsible for their production. A common feature of all the copies of TEs that generate small RNAs in the Vv3 and Sl3 strains is that they show a relatively high GC content. By contrast, the AT-rich copies of the same subfamilies do not produce small RNAs. These data corroborate those of Porquier et al. (2021), who observed that the production of small RNAs in the B05.10 strain correlates positively with the retrotransposon GC content and negatively with a RIP index. This suggests that only retrotransposons that are not inactivated by RIP are able to be expressed and therefore to generate small RNAs. Furthermore, the small RNAs generated by the seven retrotransposons present in Sl3 and/or Vv3 strains all show the same features, *i*.*e*. a size of 21-22 nucleotides and a preference for a 5′ terminal U. These features are believed to be required to associate with the plant AGO1 and activate the gene silencing process (Weiberg et al., 2013).

Among the seven small RNA generating-retrotransposons that were identified, three Gypsy subfamilies *i*.*e. Boty*/Gypsy_1, Gypsy_3 and Gypsy_4 are present in both Vv3 and Sl3 strains as well as in B05.10 (Porquier et al., 2021) and are responsible for the synthesis of a common set of small RNAs that includes the characterized effectors siR5, siR3.1 and siR3.2 (Weiberg et al., 2013). In addition to the shared repertoire of small RNAs mentioned above, the Vv3 and Sl3 strains produce additional small RNAs through the Copia_4, Gypsy_6, Gypsy_7 (Vv3) and Copia_6 (Sl3) retrotransposons. These newly characterized TEs and the small RNAs that they generate therefore confirm that the difference in the repertoires of retrotransposons present in *B. cinerea* strains Sl3 and Vv3 has strong impact on the production of potential small RNA effectors.

### The *B. cinerea* G1 population specialized on grapevine produces a specific set of small RNAs

Investigating the repertoires of small RNAs and the presence of some of the small RNA-producing TEs in a larger set of strains *B. cinerea* highlighted some significative differences between the three considered populations. While some sets of TE-derived small RNAs are present in strains belonging to different populations, some seem to be specific to one genetic group. Small RNAs produced by *Boty*/Gypsy_1, Gypsy_3 and Gypsy_4 are not only produced by the B05.10, Sl3 and Vv3 strains but appear to be commonly produced in the T and G1 populations which could suggest that this set plays a conserved role that is important in the interaction with different hosts. This is indeed the case for the characterized Gypsy_3- derived siRNAs that were shown to silence genes both in *Arabidopsis thaliana* and in tomato (Weiberg et al., 2013 ; Porquier et al., 2021). Regarding the three analyzed strains of the G2 population, only *Boty*/Gypsy_1- and Gypsy_4-derived small RNAs could be identified but uncharacterized TEs may generate other small RNAs. A full assembly of the genome of one the G2 strains would be required to answer this question.

Finally, our data revealed that two retrotransposons, Copia_4, and to a lesser extent Gypsy_7, occur at high frequency in the G1 population where they jointly generate an important proportion of the arsenal of TE-derived small RNAs. The genome of the Vv3 strain harbors ten complete copies of Copia_4 on CCs, and 23 complete copies of Gypsy_7 shared between the CCs and the AC BCIN19 that is specific to the G1 population. By contrast, these two TEs were very rarely detected by PCR in other strains of the T and G2 populations (this study) and they were not either retrieved in the genomic sequences of *B. cinerea* strains B05.10 (van Kan et al., 2017), T4 (Amselem et al., 2011) and BcDW1 (Blanco-Ulate et al., 2013). Using a larger number of strains from the G1 and G2 populations as well as strains from other hosts would be valuable to investigate further the distribution of these sources of small RNAs. If they are confirmed to be mainly present in the G1 population specialized on grapevine, it will be tempting to hypothesize that they have been maintained because they have a significant role in this interaction, either directly with the host *i*.*e*. through the silencing of grapevine genes, either indirectly *e*.*g*. by acting on other microorganisms present in the same ecological niche. Nevertheless, the fact that the G2 strains do not have the Copia_4 and Gypsy_7 TEs suggests that their potential role is not needed in all *B. cinerea* populations found on grapevine. Functional studies will be required to answer these questions.

## Conclusion

Our study was conducted in order to investigate the genomic determinants of host specialization in *B. cinerea*. Previous work revealed widespread signatures of positive selection in the T population specialized to tomato, with genes under positive selection encoding cellulases, pectinases and enzymes involved in the oxidative stress response suggesting that these activities may contribute to the specialization on tomato (Mercier et al., 2021). The present study substantially extends these findings by revealing that populations of *B. cinerea* specialized on different hosts harbor different sets of accessory chromosomes, repertoires of transposons and their derived small RNAs. Hence, we identified genomic features that are specific to the main population specialized to grapevine in France (the G1 population). The retrotransposons Copia_4 and Gypsy_7 allow the production of a G1-specific set of small RNAs with structures similar to the known effector siRNA (Weiberg et al., 2013). In addition, strains of the G1 population specifically carry the newly characterized BCIN19 accessory chromosome that includes several copies of the Gypsy_7 elements and newly identified genes whose functions remained to be characterized. Our characterization and analysis of new genomic data in populations of *Botrytis* specialized to different hosts pave the way for new molecular investigations of the mechanisms underlying host specialization in this polyphagous pathogen.

## Material and methods

### Genome sequencing and assembly

The *Botrytis cinerea* Sl3 and Vv3 strains were isolated from the Champagne region (France) respectively from tomato (cultivar Moneymaker) and grapevine berries (Pinot noir), as previously described (Mercier et al., 2021). For DNA sequencing, these strains were cultivated three days in liquid malt medium. High-molecular-weight DNA was extracted using a sarkosyl procedure in which the ethanol-precipitated DNA was fished with a glass hook to avoid centrifugation and DNA breaks. Sl3 and Vv3 genomic DNA were sequenced using a PacBio Sequel sequencer (KeyGene, Wageningen, NL). Total lengths of respectively 8.3Gb and 4.7Gb reads were obtained with mean lengths of 8.4Kb and 11.3Kb. The theorical coverages of the genome were thus of 195X and 120X. Reads were assembled with HGAP4 (SMRT-Link v5.0.1) (https://github.com/PacificBiosciences/) and Canu v1.6 (Koren et al., 2017). HGAP4 was run with the set of parameters recommended for fungal genomes (https://pb-falcon.readthedocs.io/en/latest/parameters.html). Canu was run with default parameters and an expected coverage of 42Mb. HGAP assemblies were polished with pilon (Walker et al., 2014) using Sl3 or Vv3 Illumina reads (Mercier et al., 2021). Canu assemblies were polished with both arrow (https://github.com/PacificBiosciences/) and PILON. Mitochondrial, ribosomal and small redundant contigs were removed, and final versions of assemblies were manually compiled from polished runs of HGAP and Canu assemblies. Both genome assemblies were evaluated and compared with B05.10 referent genome (van Kan et al., 2017) using Quast (Gurevich et al., 2013).

For the separation of ACs by gel electrophoresis, chromosomal DNA was prepared as described by van Kan et al. (1993) and loaded on a 1% agarose gel (SeaKem Le, FMC) in 0.5X Tris Borate EDTA buffer. Chromosomes were then separated using a Contour-clamped Homogeneous Electric Field (CHEF) apparatus (DRIII, BioRad) using the following parameter: 6 V/cm; 120° angle; 50-90 seconds switch; 22 hours; 14°C.

### Transposable elements prediction

Transposable elements (TEs) were annotated using the REPET package (https://urgi.versailles.inra.fr/Tools/REPET ; (Amselem et al., 2015) as previously described in Porquier et al., 2016 for the genome of the *B. cinerea* reference strain B05.10. Briefly, the TEdenovo pipeline (Flutre et al., 2011) was used to detect repeated elements in the genome and to provide a consensus sequence for each family. Consensus sequences were then classified using the PASTEC tool (Hoede et al., 2014), based on the Wicker hierarchical TE classification system (Wicker et al., 2007). After manual correction, the resulting library of consensus sequences was used to annotate TE copies in the whole genome using the TEannot pipeline (Quesneville et al., 2005). Consensus sequences identified from different genomes were compared and considered as the same subfamily based on a bidirectional best hit approach (blastn with evalue < 1e-10 and coverage > 70%). Presence of the subfamilies in the genomes of additional strains (Table 1) was tested by PCR using the MyTaq polymerase (Bioline) and the primers listed in Sup. Table S5.

### Gene prediction, synteny

The structural annotation of B05.10 genes (van Kan et al., 2017) (http://fungi.ensembl.org/Botrytis_cinerea/) was transferred to Vv3 and Sl3 genomes using the Liftoff annotation mapping tool (Shumate & Salzberg, 2021). Genes were also predicted *de novo* using the Fgenesh *ab initio* gene-finder (Solovyev et al., 2006) (http://www.softberry.com/berry.phtml) with the *Botrytis* matrix. Presence/absence of genes in other genomes was determined with blastn analyses. The synteny between B05.10, Sl3 and Vv3 genes was analysed with SynChro (Drillon et al., 2014), which detects ortholog proteins with Reciprocal Best Hit (RBH). SynChro was run with a best score threshold (min_sim_RBH) of 80, a length ratio threshold (max_diflen) of 1.3 and a delta value of 3 (medium stringency). Duplications of gene clusters were further explored with CLINKER (Gilchrist & Chooi, 2021). Presence of the gene *Bctlh* in the genomes of additional strains (Table 1) was tested by PCR using the MyTaq polymerase and the primers indicated in Sup. Table S5.

### Small RNA sequencing and analysis

*Botrytis cinerea* strains were cultivated 48 hours on solid media made from grape juice or tomato juice supplemented with agar and covered with a sheet of cellophane as previously described (Simon et al., 2013). RNAs were extracted with the TRIzol reagent (InvitroGen) and submitted to a DNAse treatment (DNA-free kit, Ambion). Total RNAs were purified with miRNeasy kit (Qiagen) which allows the selection of the RNA fraction less than 100 bases. Small RNA libraries were prepared and sequenced at Integragen (https://www.integragen.com/). Libraries were generated following (Vigneault et al., 2012), with adjustments to improve ligation, from at least 1 μg of extracted total RNA with a RIN greater than 7. Libraries were sequenced on Illumina HiSeq4000. Quality of raw sequence data was checked with FastQC (https://www.bioinformatics.babraham.ac.uk/projects/fastqc/). After removal of adapters with Cutadapt (Martin, 2011), reads shorter than 16 bases were discarded. Reads were then processed using FASTX-TOOLKIT (http://hannonlab.cshl.edu/fastx_toolkit/) as in Weiberg et al. (2013), *i*.*e*. low-quality sequences were filtered out with FASTQ_QUALITY_FILTER -q 30 -p 70, low-complexity sequences were filtered out with FASTX_ARTIFACTS_FILTER, and finally identical sequences were counted with FASTX_COLLAPSER. Only sequences 20-24 nucleotides length and with more than five reads per million in at least one library were kept for further analyses. Principal Component Analyses (PCA) were performed to visualize the distance between samples. Reads were mapped against B05.10 (van Kan et al., 2017), Sl3 and Vv3 genomes, against TE consensus and against TE complete copies in the three genomes with the GLINT software (https://forge-dga.jouy.inra.fr/projects/glint/wiki); only perfect matches (100% identity and 100% coverage) were considered. As a control, unprocessed (raw) reads were mapped the same way to verify that no additional small RNA-producing TE could be identified.

## Acknowledgements

We are grateful to the following bioinformatics platforms and partners for providing computational support and/or storage resources: Bioinfo Genotoul, (https://doi.org/10.15454/1.5572369328961167E12), CATI BARIC (https://www.cesgo.org/catibaric/), INRAE-URGI, INRAE-LIPME Bioinfo (Jérôme Gouzy, Sébastien Carrère, Erika Sallet) and INRAE-BIOGER Bioinfo (https://bioinfo.bioger.inrae.fr/ ; Nicolas Lapalu). We thank Alexander H. J. Wittenberg (KeyGene, Wageningen, NL) and Jean-Paul Saraiva (Integragen, Evry, France) for the PacBio and Illumina sequencing, respectively. Preprint version 4 of this article has been peer-reviewed and recommended by Peer Community In Genomics (https://doi.org/10.24072/pci.genomics.100023).

## Availability of data and materials

The genomic raw data were deposited into the NCBI SRA under the accessions PRJNA752967 for Sl3 and PRJNA752962 for Vv3. Genome assemblies were deposited at NCBI under the accessions CP080979-CP080996 for Sl3 and CP080961-CP080978 for Vv3. Small RNAs raw data and the matrix of sequence counts were deposited into the NCBI GEO under accession GSE181592. Furthermore, the genomic and annotation (gene/TE) fasta and gff files, as well as Sl3 and Vv3 genome browsers are available at the INRAE BIOGER Bioinformatics platform (https://bioinfo.bioger.inrae.fr/).

## Supplemental information

Supplemental information is available at https://doi.org/10.1101/2022.03.07.483234

**File S1**: Consensus sequences of the repeated elements identified in the genomes of *B. cinerea* strains Sl3 and Vv3.

**Table S1:** Chromosomes of the *B. cinerea* strains B05.10, Sl3 and Vv3.

**Table S2:** Genes predicted in the genomes of *B. cinerea* strains B05.10, Sl3 and Vv3.

**Table S3**: Repertoires of Transposable Elements (TEs) in the *B. cinerea* strains B05.10, Sl3 and Vv3.

**Table S4**: Libraries of small RNAs isolated from *B. cinerea* strains belonging to different populations.

**Table S5:** List of PCR primers used to detect the presence of Transposable Elements (TEs) or the gene encoding the Telomere-Linked Telomerase (*BcTLH*).

**Figure S1:** Genomic genealogy of *B. cinerea* strains showing the populations T, G1 and G2.

**Figure S2:** CHEF gel electrophoresis resolving accessory chromosomes of *B. cinerea* strains B05.10, Sl3 and Vv3.

**Figure S3:** Synteny between the Core Chromosomes (CCs) of the *B. cinerea* strains B05.10, Sl3 and Vv3.

**Figure S4**: Main inversions events observed between the genomes of the *B. cinerea* strains Sl3 and Vv3.

**Figure S5:** Localization of the PKS7 secondary metabolism gene cluster in the genomes of the *B. cinerea* strains B05.10 and Vv3.

**Figure S6**: Duplication of four contiguous genes in the genome of the *B. cinerea* strain Sl3.

**Figure S7:** Structure of the Telomere-Linked Helicase (TLH) identified in the *B. cinerea* strain Vv3.

**Figure S8:** Positions of the genes encoding the Telomere-Linked Helicase (BcTLH) and of the Helitron-like Transposable Elements (TE) in the genome of the *B. cinerea* strain Vv3.

**Figure S9:** Alignment of the ten consensus sequences corresponding to the *Boty*/Gypsy_1 Transposable Element (TE) identified in the genomes of the *B. cinerea* strains B05.10, Sl3 or Vv3.

**Figure S10:** Helitron-like Transposable Element (TE) identified in the genome of the *B. cinerea* strain Vv3.

**Figure S11:** Localization of the main superfamilies of Transposable Elements (TEs) in the genomes of the *B. cinerea* strains Sl3 and Vv3.

**Figure S12:** GC content of complete copies of the main subfamilies of Transposable Elements (TEs) in the genomes of the *B. cinerea* strains B05.10, Sl3 and Vv3.

**Figure S13:** Principal Component Analysis (PCA) of the repertoires of small RNAs in different strains of *B. cinerea*.

**Figure S14:** Retrotransposons that generate small RNAs in the *B. cinerea* strains Sl3 and/or Vv3.

**Figure S15:** Mapping of small RNA reads on complete copies of TEs in relation to their GC percent.

**Figure S16:** Small RNAs produced by the *B. cinerea* strains Sl3 and Vv3: size distribution and 5’ nucleotide.

**Figure S17:** Positions of complete copies of the TE Copia_4, Gypsy_6 and Gypsy_7 in the *B. cinerea* Vv3 genome.

**Figure S18:** Alignment of Copia_4, Gypsy_6 and Gypsy_7 consensus sequences with their respective complete copies in Vv3 genome.

