## Supplemental Information for "*Botrytis cinerea* strains infecting grapevine and tomato display contrasted repertoires of accessory chromosomes, transposons and small RNAs"

### Supplemental file:

**File S1: Consensus sequences of the repeated elements identified in the genomes of *B. cinerea* strains Sl3 and Vv3.** The consensus were determined and annotated with the REPET pipeline (Flutre et al., 2011; Amselem et al., 2015).

(Fasta file included separately)

### Supplemental tables:

**Table S1: Chromosomes of the *B. cinerea* strains B05.10, Sl3 and Vv3.** B05.10 data are those of van Kan et al. (2017). The percentage of B05.10 chromosomes coverage was calculated with QUAST (Gurevich at al., 2013) with the default parameters. The presence of telomeric repeats (*i.e.* TTAGGG repetitions) was sought at the extremities of chromosomes (ν indicates the presence). Genes previously identified in B05.10 were transferred with LIFTOFF (Shumate & Salzberg, 2021) and a *de novo* gene prediction was performed (FGENESH, Solovyev et al., 2006) to identify additional genes. See **Supplementary Table S2** for details.

(Excel file included separately)

**Table S2: Genes predicted in the genomes of *B. cinerea* strains B05.10, Sl3 and Vv3.** B05.10 data are those of van Kan et al. (2017). Genes previously identified in B05.10 were transferred with LIFTOFF (Shumate & Salzberg, 2021) and a *de novo* gene prediction was performed (FGENESH, Solovyev et al., 2006) to identify additional genes.

(Excel file included separately)

**Table S3: Repertoires of Transposable Elements (TEs) in the *B. cinerea* strains B05.10, Sl3 and Vv3.** Consensus sequences of the subfamilies of TEs identified in Sl3 and Vv3 genomes were compared to those previously identified in B05.10 and numbered accordingly (Porquier et al., 2016; Porquier et al., 2021). Percentages in boxes represent the coverage of the subfamily in the corresponding genome, while white boxes represent absent subfamilies. The number of Full-Length Copies (FLC) and the resulting coverage were also computed.

| | | | | | | | | | Total coverage | | | # fullLgthCopies (FLC) | | | coverage by FLC | | |
|---|---|---|---|---|---|---|---|---|---|---|---|---|---|---|---|---|---|
| Class | Order | Superfamily | W_CODE | *Subfamily name / consensus name* | | *Len (Kb)* | *Consensus name in B05.10 (Porquier et al., 2016)* | *Subfamily name in B05.10 (Porquier et al., 2021)* | *B0510* | *Sl3* | *Vv3* | *B0510* | *Sl3* | *Vv3* | *B0510* | *Sl3* | *Vv3* |
| I | LTR | Copia | RLC | Copia_1 | | 6.4 | RLX_P27.0 | BcCopia1 | 0.25% | 0.39% | 0.54% | 12 | 22 | 29 | 0.18% | 0.33% | 0.41% |
| | | | | Copia_2 | | 8.2 | RLX_G54 | BcCopia2 | 0.14% | 0.12% | 0.16% | 6 | 4 | 5 | 0.12% | 0.08% | 0.09% |
| | | | | Copia_3 | | 5.4 | | | | | 0.18% | | | 14 | | | 0.17% |
| | | | | Copia_4 | | 5.4 | | | | | 0.14% | | | 10 | | | 0.12% |
| | | | | Copia_5 | | 6 | | | | | 0.63% | | | 37 | | | 0.50% |
| | | | | Copia_6 | | 5.5 | | | | 0.14% | | | 8 | | | 0.10% | |
| | | Gypsy | RLG | Boty/Gypsy_1 | Boty-var1 | 6.6 | RLX_P26.1 | BcGypsy1 | 0.27% | 0.30% | 0.77% | 13 | 15 | 25 | 0.20% | 0.23% | 0.38% |
| | | | | | Boty-var2 | 6.5 | RLX_P21.1 | | 0.43% | 0.17% | 0.15% | 21 | 7 | 9 | 0.33% | 0.11% | 0.12% |
| | | | | | Boty-var3 | 6.4 | RLX_P6.1 | | 0.14% | 0.18% | 0.18% | 7 | 10 | 8 | 0.11% | 0.15% | 0.11% |
| | | | | | Boty-var4 | 6.5 | RLX_P13.1 | | 0.12% | 0.12% | 0.07% | 7 | 6 | 3 | 0.11% | 0.09% | 0.04% |
| | | | | | Boty-var5 | 6.1 | | | | | 0.26% | | | 18 | | | 0.25% |
| | | | | | Boty-var6 | 6.3 | | | | | 0.15% | | | 7 | | | 0.10% |
| | | | | | Boty-var7 | 6.2 | | | | | 0.09% | | | 5 | | | 0.07% |
| | | | | | Boty-var8 | 6.2 | | | | | 0.07% | | | 5 | | | 0.07% |
| | | | | | Boty-var9 | 6.6 | | | | 0.25% | | | 14 | | | 0.21% | |
| | | | | | Boty-var10 | 6.6 | | | | 0.27% | | | 11 | | | 0.17% | |
| | | | | Gypsy_2 | | 6.4 | RLX_P17.7 | BcGypsy2 | 0.14% | 0.12% | 0.18% | 8 | 6 | 9 | 0.12% | 0.09% | 0.13% |
| | | | | Gypsy_3 | | 7.4 | RLX_B-R56 | BcGypsy3 | 0.28% | 0.35% | 0.34% | 13 | 20 | 11 | 0.23% | 0.34% | 0.18% |
| | | | | Gypsy_4 | | 11 | RLX_G57 | BcGypsy4 | 0.57% | 0.50% | 0.39% | 13 | 14 | 6 | 0.33% | 0.36% | 0.15% |
| | | | | Gypsy_5 | | 6.5 | | | | | 0.11% | | | 6 | | | 0.09% |
| | | | | Gypsy_6 | | 6.7 | | | | | 0.21% | | | 12 | | | 0.18% |
| | | | | Gypsy_7 | | 6.3 | | | | | 0.34% | | | 23 | | | 0.32% |
| | TRIM | TRIM | RXX-TRIM | Trim_1 | | 0.5 | | | | | 0.01% | | | 7 | | | 0.01% |
| | | | | Trim_2 | | 0.5 | | | | | 0.01% | | | 5 | | | 0.01% |
| II | TIR | Tc1-Mariner | DTT | Flipper | | 1.8 | DTX_P14.9 | | 0.06% | 0.14% | 0.19% | 10 | 29 | 41 | 0.04% | 0.12% | 0.17% |
| | | | | Mariner_1 | | 1.9 | DTX_G36 | | 0.09% | 0.13% | 0.13% | 16 | 25 | 26 | 0.07% | 0.11% | 0.11% |
| | | | | Mariner_2 | | 1.9 | | | | 0.38% | 0.26% | | 54 | 29 | | 0.23% | 0.12% |
| | | | | Mariner_3 | | 1.9 | | | | 0.15% | 0.12% | | 21 | 16 | | 0.09% | 0.07% |
| | | | | Mariner_4 | | 1.9 | | | | | 0.11% | | | 18 | | | 0.08% |
| | | | | Mariner_5 | | 1.9 | | | | | 0.08% | | | 27 | | | 0.11% |
| | | | DTX | TIRnoCat | | 0.7 | | | | | 0.15% | | | 51 | | | 0.08% |
| | Helitron | Helitron | DHH | Helitron | | 15.5 | | | | | 1.37% | | | 8 | | | 0.28% |
| | | | DXX-MITE | MITE_1 | | 0.5 | DXX-MITE_G19 | | 0.04% | | 0.02% | 6 | | 15 | 0.01% | | 0.02% |
| | | | | MITE_2 | | 0.5 | DXX-MITE_P15.8 | | 0.47% | | | 92 | | | 0.10% | | |
| | | | | MITE_3 | | 0.4 | | | | | 0.23% | | | 98 | | | 0.10% |
| | | | | MITE_4 | | 0.5 | | | | | 0.03% | | | 15 | | | 0.02% |
| | | | | MITE_5 | | 0.5 | | | | 0.02% | | | 3 | | | 0.00% | |
| noCat | | | | noCat_1 | | 0.6 | noCat_G9 | | 0.45% | | | 38 | | | 0.05% | | |
| | | | | noCat_2 | | 13.9 | noCat_R20 | | 0.27% | | | 1 | | | 0.03% | | |
| | | | | noCat_3 | | 0.5 | | | | | 0.02% | | | 8 | | | |
| | | | | noCat_4 | | 19.7 | | | | 0.40% | | | 6 | | | 0.27% | |
| | | | | noCat_5 | | 6.8 | | | | 0.40% | | | 22 | | | 0.35% | |
| **Coverage by shared subfamilies (B05.10, Sl3, Vv3)** | | | | | | | | | **2.49%** | **3.04%** | **3.65%** | | | | **1.84%** | **2.38%** | **2.38%** |
| **Coverage by specific subfamilies (only one genome)** | | | | | | | | | **1.19%** | **0.96%** | **3.62%** | | | | **0.18%** | **0.73%** | **2.04%** |
| **Coverage by subfamilies shared by 2 genomes** | | | | | | | | | **0.04%** | **0.53%** | **0.41%** | | | | **0.01%** | **0.33%** | **0.21%** |
| **Nb consensus** | | | | 42 consensus (33 subfamilies) | | | | | 15 | 19 | 33 | | | | | | |
| **Genome coverage** | | | | | | | | | **3.72%** | **4.53%** | **7.70%** | | | | **2.02%** | **3.44%** | **4.53%** |

**Table S4: Libraries of small RNAs isolated from *B. cinerea* strains belonging to different populations**. Strains were grown either on grape or tomato juice medium (GJ and TJ, respectively). The number of reads after each successive quality filtering step is indicated. Finally, only sequences of length 20-24nt and with at least five reads per million in at least one library were mapped on the Transposable Elements (TEs) identified in the strains B05.10, Sl3 and Vv3. The Reads Per Million (RPMs) values above 300 are reported for each TE consensus.

(Excel file included separately)

**Table S5: List of PCR primers used to detect the presence of Transposable Elements (TEs) or the gene encoding the Telomere-Linked Telomerase (*BcTLH*).**

| TE or gene | Primer forward | | Primer Reverse | | Size of the PCR product |
|---|---|---|---|---|---|
| Helitron-like TE | MV55 | TGATCGATTCGCTGACCTCG | MV56 | CGGAGTTATGAGACCGGACG | 262 bp |
| BcTLH | MV148 | GTGGGGTTTGATGCATGTGT | MV149 | GCAATCCTTGTAACCGGTCC | 1292 bp |
| Copia 4 TE | AS934 | ACACTATAGAGCCCAGCGAC | AS933 | CCAATGAGAAGGGGTGGTCT | 205 bp |
| Gypsy 6 TE | MV117 | TGATACCACGCTCCTTACCC | MV118 | TCGGTCTTCTGAGGCTTTGT | 340 bp |
| Gypsy 7 TE | MV115 | CTACGGAAGCCACTATACAC | MV116 | ACTTGTACAGCGGCTTCGAT | 363 bp |

### Supplemental figures:

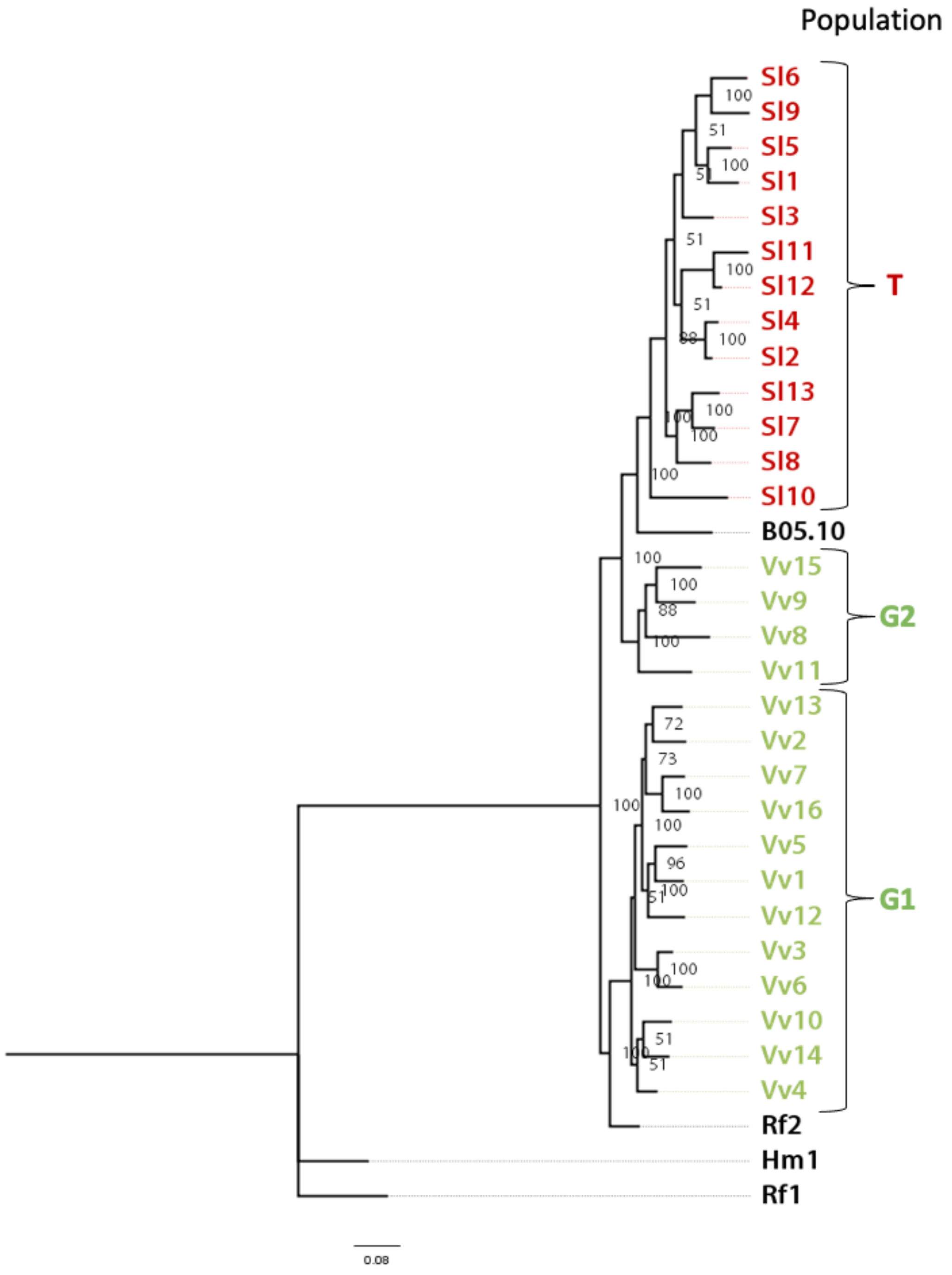


**Figure S1: Genomic genealogy of *B. cinerea* strains showing the populations T, G1 and G2.** This Maximum likelihood tree was generated from our previous genomic data (Mercier et al., 2021) using the RaxML (Stamatakis, 2014) software with GTR-CAT model of nucleotide substitution under a Gamma model of rate heterogeneity with 1000 bootstrap replicates. Node labels represent bootstrap support. The tree was built upon the 249,084 filtered SNPs (*i.e.* monomorphic sites were excluded), so branch lengths are expressed in terms of variant sites. Strains sampled from *Vitis vinifera* (Vv) are colored in green, those from *Solanum lycopersicum* (Sl) are in red and those from other hosts (Rf, *Rubus fruticosus*; Hm, *Hydrangea macrophylla*) are in black.

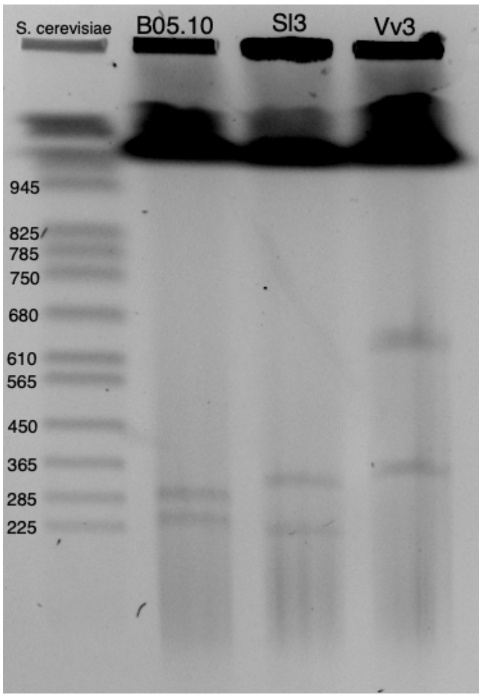


**Figure S2: CHEF gel electrophoresis resolving smaller chromosomes of *B. cinerea* strains B05.10, Sl3 and Vv3.** The size of the chromosomes of *Saccharomyces cerevisiae* are indicated on the left (Kb).

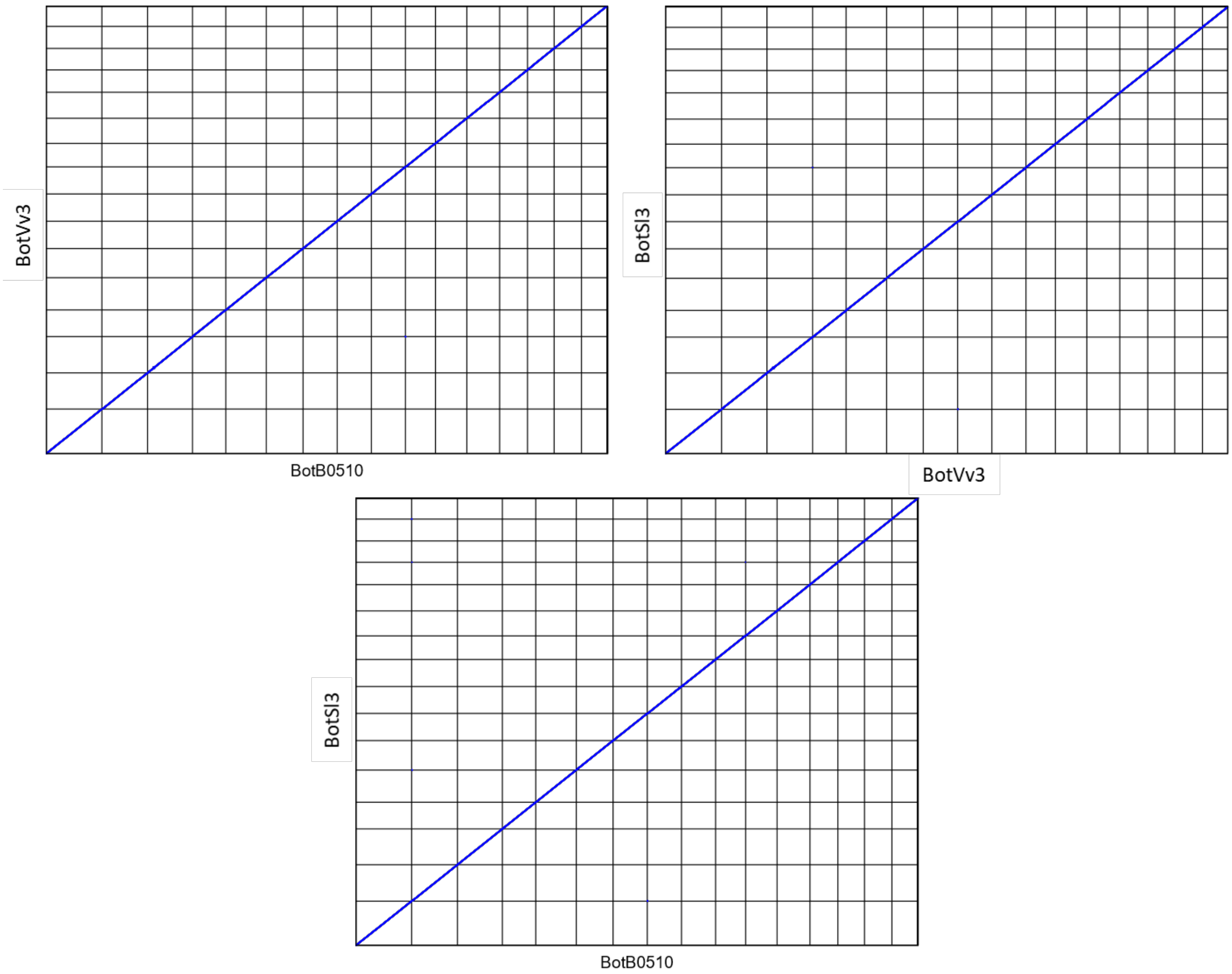


**Figure S3: Synteny between the Core Chromosomes (CCs) of the *B. cinerea* strains B05.10, Sl3 and Vv3**. DotPlots of the genes detected on CCs 1 to 16 were generated with SynChro (Drillon et al., 2014).

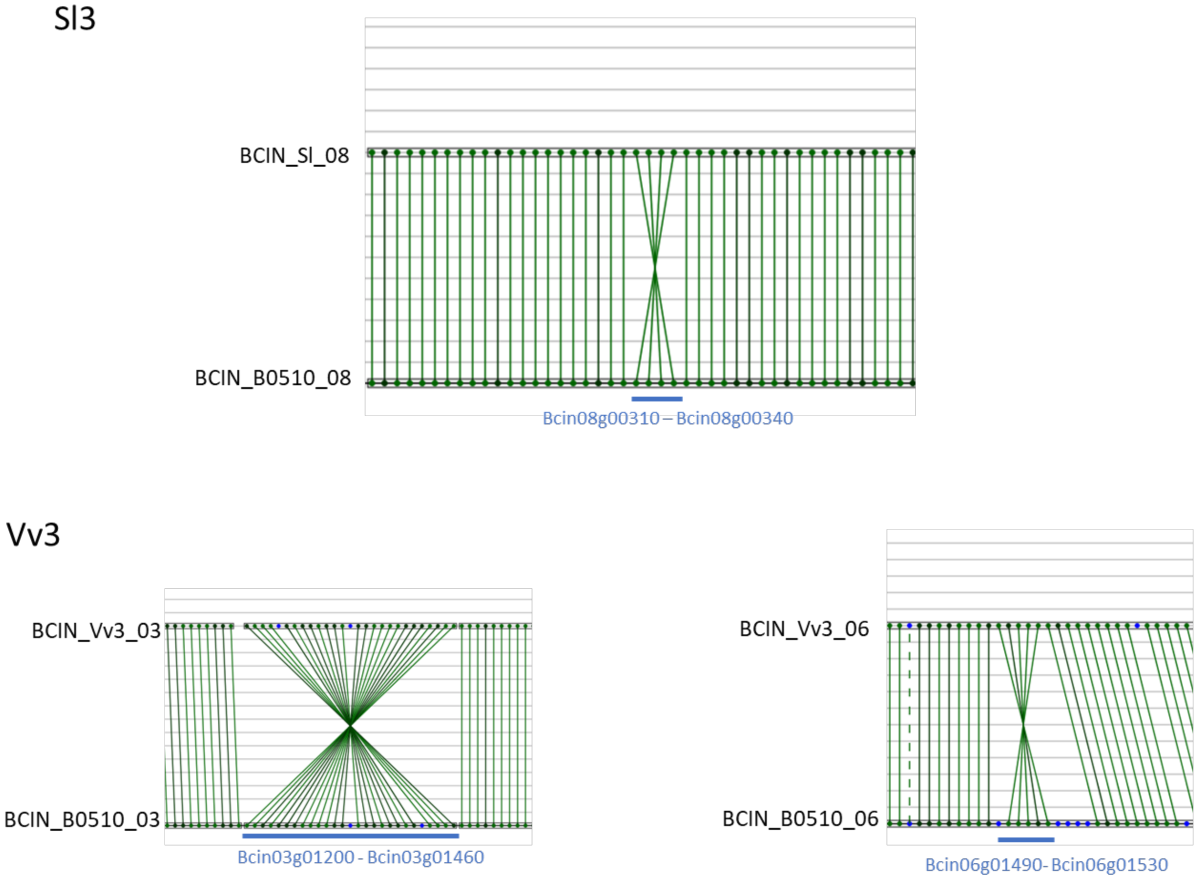


**Figure S4: Main inversions events observed between the genomes of the *B. cinerea* strains Sl3 and Vv3.** The figures were obtained thank to SYNCHRO (Drillon et al., 2014) with the genome of the strain B05.10 as a reference. Dots symbolize genes, plain lines highlight RBH (Reciprocal Best Hits) relationships and dotted lines represent non-RBH homologous relationships.

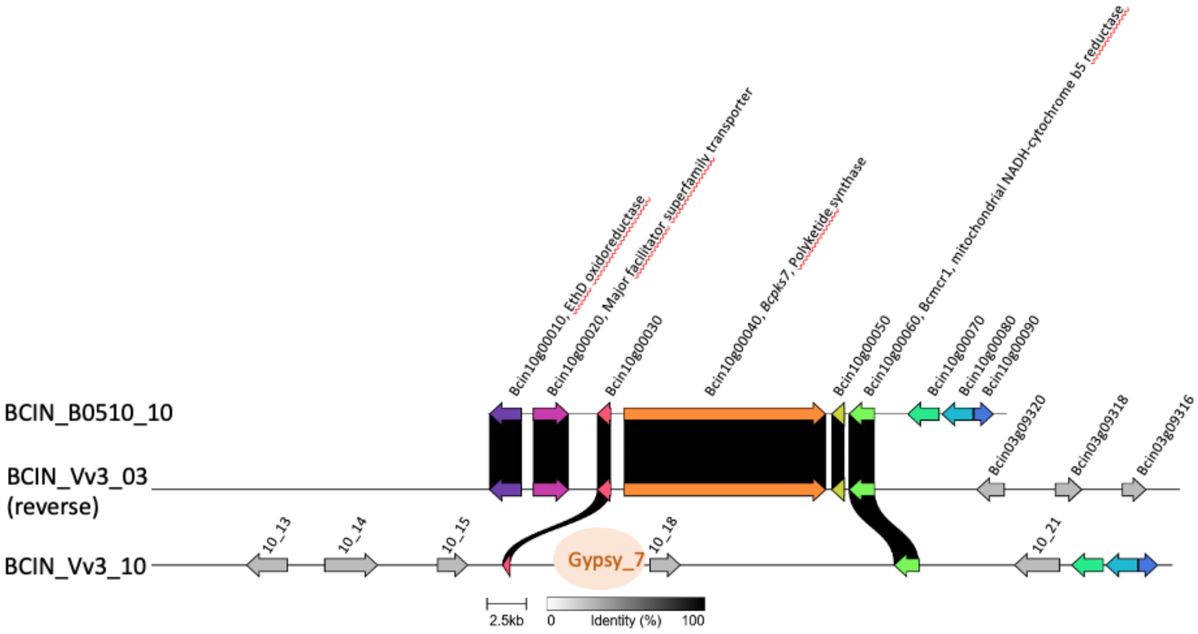


**Figure S5: Localization of the PKS7 secondary metabolism gene cluster in the genomes of the *B. cinerea* strains B05.10 and Vv3.** The PKS7 secondary metabolism gene cluster includes the genes Bcin10g00010 to Bcin10g00040. The left side of the figure corresponds to the extremities of the chromosomes BCIN_10 of B05.10 and of the chromosomes BCIN_Vv3_03 (in its reverse orientation) and BCIN_Vv3_10 of Vv3. The proteins were aligned and visualized with CLINKER (Gilchrist & Chooi, 2021). Gypsy_7 is a retrotransposon.

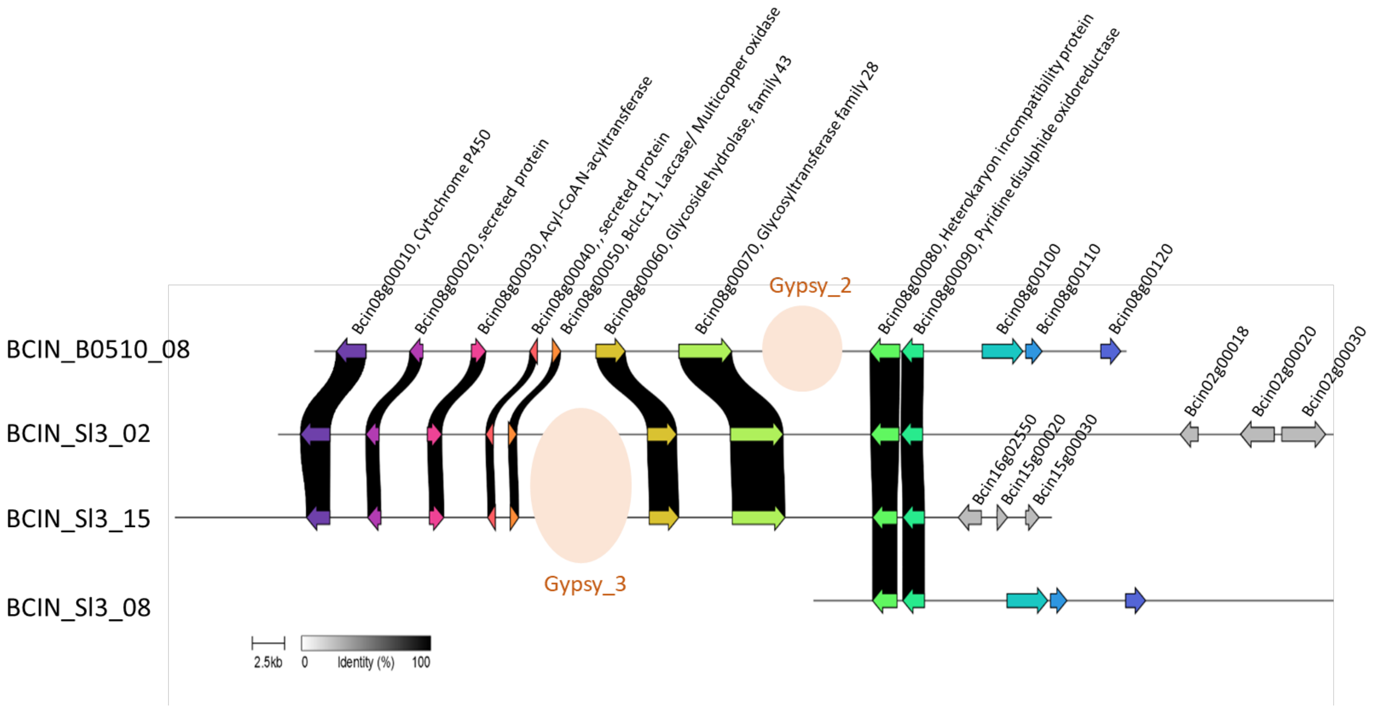


**Figure S6: Duplication of four contiguous genes in the genome of the *B. cinerea* strain Sl3**. The left side of the figure corresponds to the extremities of the chromosomes BCIN_08 of B05.10 and of the chromosomes BCIN_Sl3_02, BCIN_Sl3_15 and BCIN_Sl3_08 of Sl3. The proteins were aligned and visualized with CLINKER (Gilchrist & Chooi, 2021). Gypsy_2 and Gypsy_3 are retrotransposons.

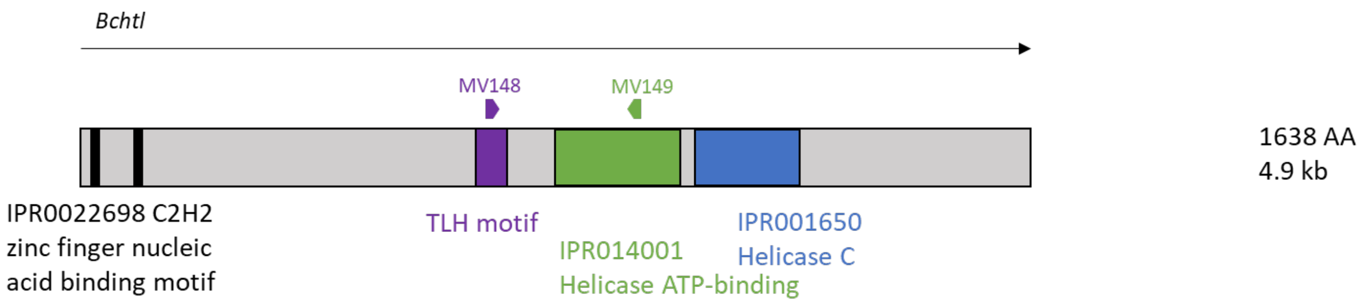


**Figure S7: Structure of the Telomere-Linked Helicase (TLH) identified in the *B. cinerea* strain Vv3**. The predicted protein BCIN_Vv3_1442 was scanned with INTERPROSCAN. Alignement with the TLHs from *Magnaporthe grisea* further allowed to identify the TLH specific motif defined by Rehmeyer et al. (2009; amino acid 625 to 790). Primers MV148 and MV149 were used to detect *Bctlh* in additional strains of *B. cinerea*.

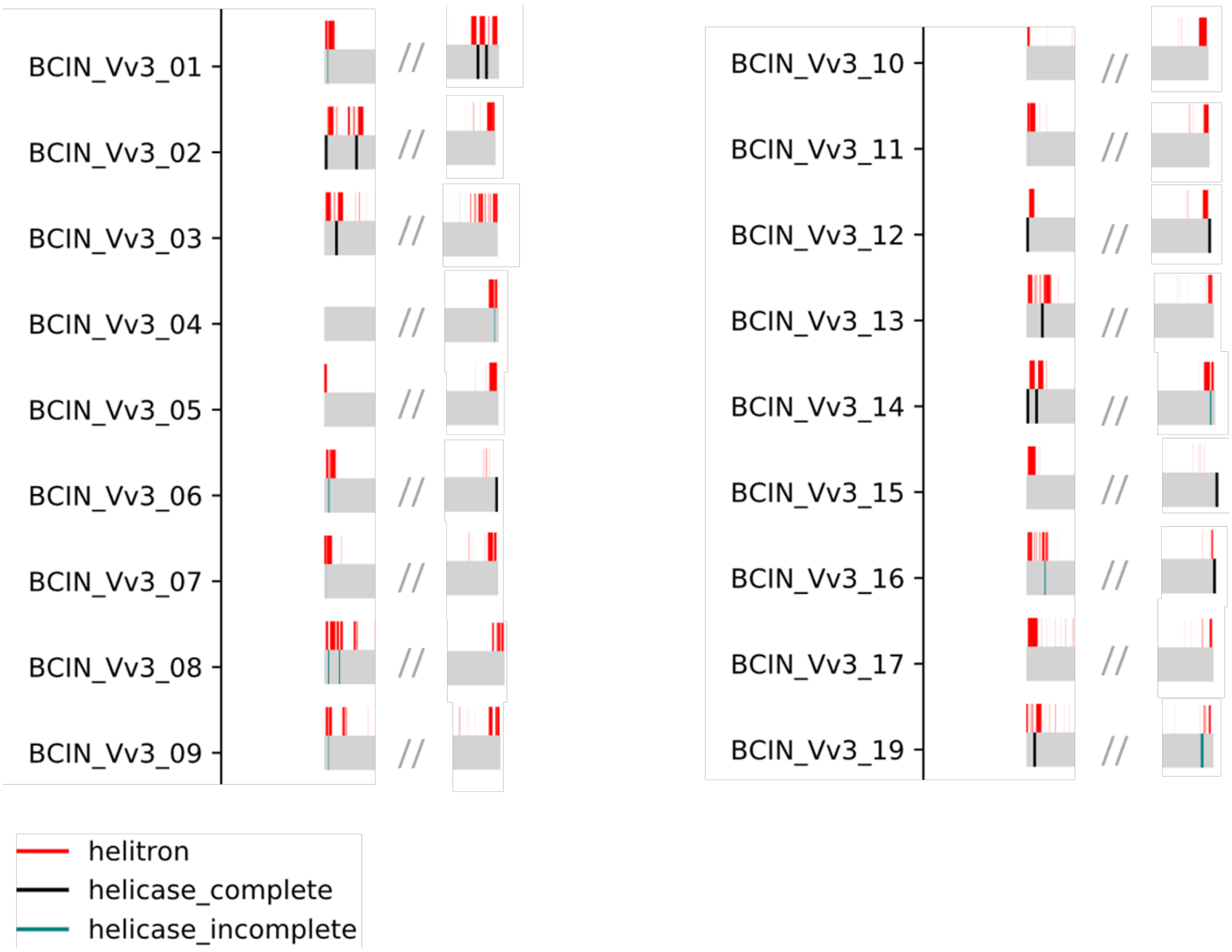


**Figure S8: Positions of the genes encoding the Telomere-Linked Helicase (BcTLH) and of the Helitron-like transposable elements in the genome of the *B. cinerea* strain Vv3**. As all the copies of *Bctlh* are located in subtelomeric regions, only these regions are displayed.

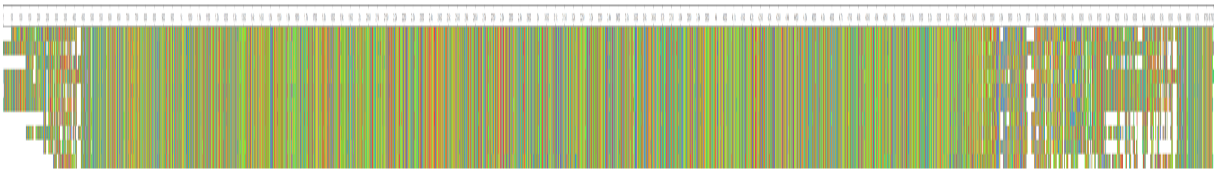

**Figure S9: Alignment of the ten consensus sequences corresponding to the Boty_Gypsy_1 transposable element identified in the genomes of the *B. cinerea* strains B05.10, Sl3 or Vv3.**

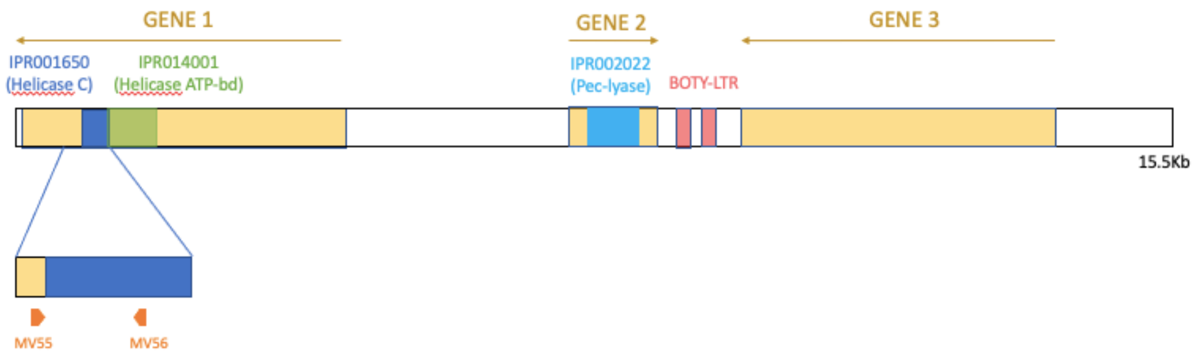


**Figure S10: Helitron-like Transposable Element (TE) identified in the genome of the *B. cinerea* strain Vv3.** Structure of the Helitron-like TE. Gene 1 encodes a protein with a helicase domain that is typical of Helitron DNA transposons, while genes 2 and 3 are probable captured genes. Gene 2 is predicted to encode a pectate lyase. Primers MV55 and MV56 were used to detect this element in additional strains of *B. cinerea*.

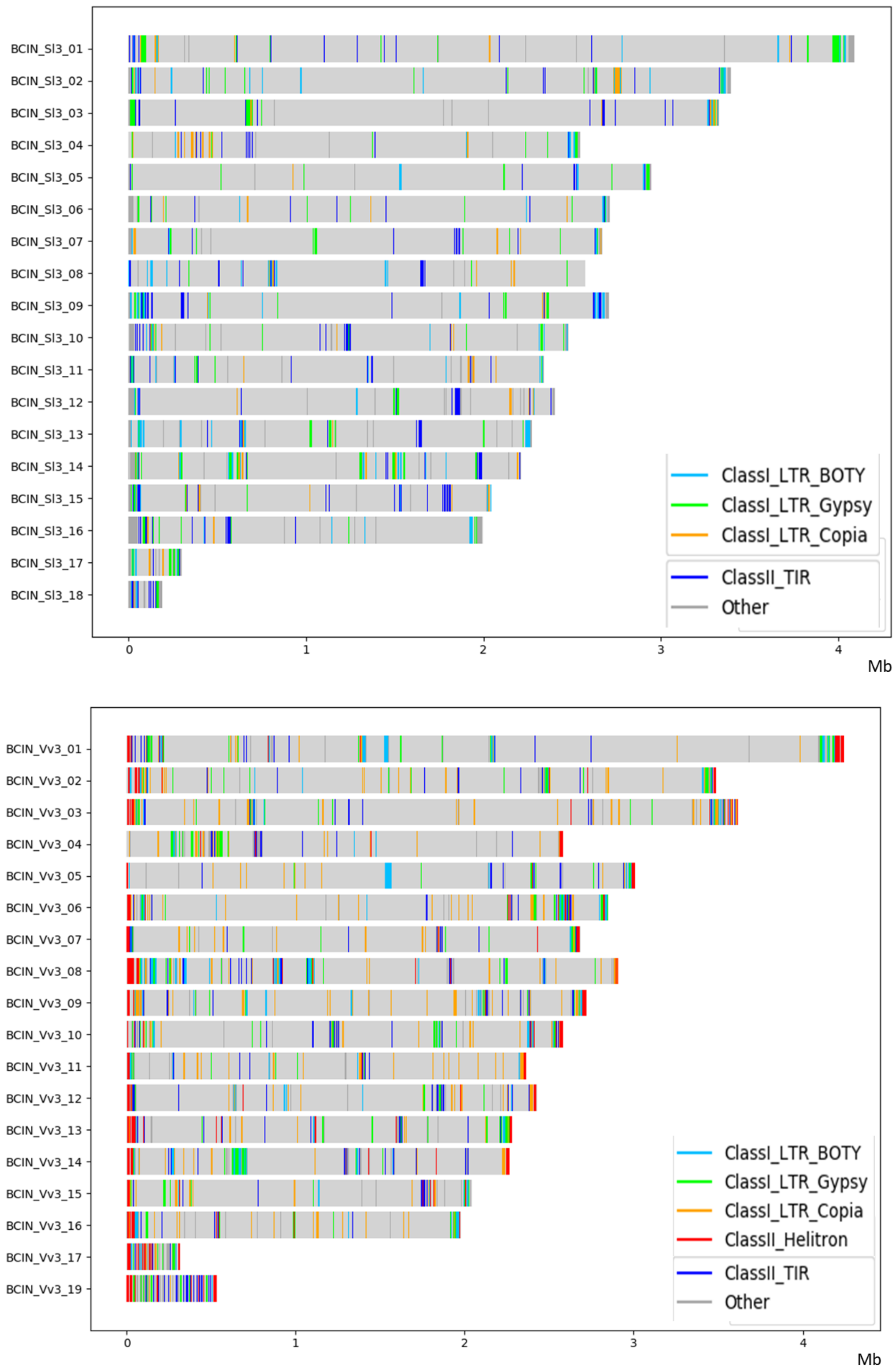


**Figure S11: Localization of the main superfamilies of Transposable Elements (TEs) in the genomes of the *B. cinerea* strains Sl3 and Vv3.**

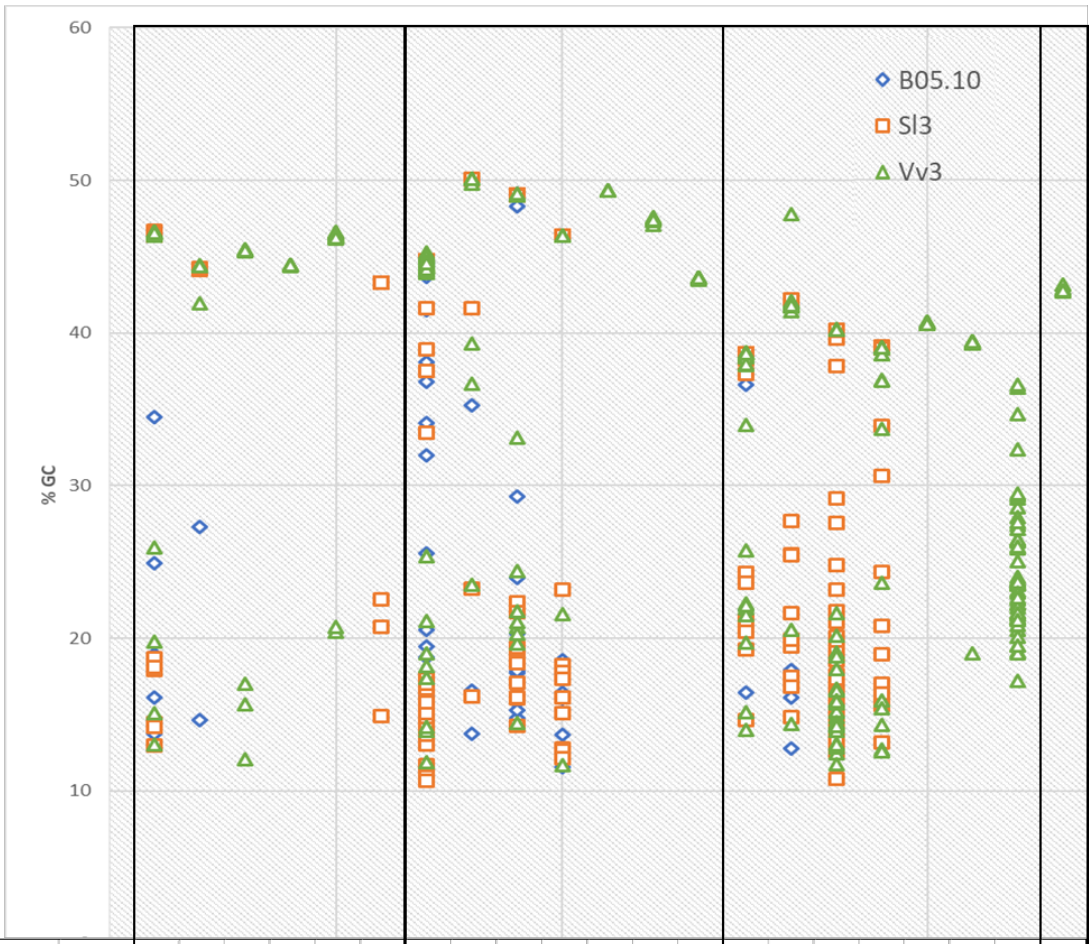


| | | Copia_1 | Copia_2 | Copia_3 | Copia_4 | Copia_5 | Copia_6 | Boty_Gypsy_1 | Gypsy_2 | Gypsy_3 | Gypsy_4 | Gypsy_5 | Gypsy_6 | Gypsy_7 | Flipper | Mariner_1 | Mariner_2 | Mariner_3 | Mariner_4 | Mariner_5 | TIR_noCat | Helitron |
|---|---|---|---|---|---|---|---|---|---|---|---|---|---|---|---|---|---|---|---|---|---|---|
| **Number of complete copies** | B0510 | **12** | **6** | | | | | **48** | **8** | **13** | **13** | | | | **10** | **16** | | | | | | |
| GC% > 40 | | 5 | 4 | | | | | 29 | 5 | 2 | 4 | | | | 0 | 12 | | | | | | |
| 30 < %GC < 40 | | 1 | 0 | | | | | 4 | 1 | 0 | 0 | | | | 5 | 0 | | | | | | |
| 20 < %GC < 30 | | 1 | 1 | | | | | 2 | 0 | 3 | 0 | | | | 2 | 1 | | | | | | |
| GC% < 20 | | 5 | 1 | | | | | 13 | 2 | 8 | 9 | | | | 3 | 3 | | | | | | |
| **Number of complete copies** | Sl3 | **22** | **4** | | | | **8** | **63** | **6** | **20** | **14** | | | | **29** | **25** | **54** | **21** | | | | |
| GC% > 40 | | 15 | 4 | | | | 5 | 44 | 4 | 10 | 4 | | | | 0 | 16 | 7 | 0 | | | | |
| 30 < %GC < 40 | | 0 | 0 | | | | 0 | 3 | 0 | 0 | 0 | | | | 22 | 0 | 2 | 14 | | | | |
| 20 < %GC < 30 | | 0 | 0 | | | | 2 | 0 | 1 | 2 | 1 | | | | 5 | 4 | 8 | 2 | | | | |
| GC% < 20 | | 7 | 0 | | | | 1 | 16 | 1 | 8 | 9 | | | | 2 | 5 | 37 | 5 | | | | |
| **Number of complete copies** | Vv3 | **29** | **5** | **14** | **10** | **37** | | **80** | **9** | **11** | **6** | **6** | **12** | **23** | **41** | **26** | **29** | **16** | **18** | **27** | **51** | **8** |
| GC% > 40 | | 25 | 5 | 11 | 10 | 35 | | 72 | 6 | 3 | 4 | 6 | 12 | 23 | 0 | 24 | 3 | 0 | 18 | 0 | 0 | 8 |
| 30 < %GC < 40 | | 0 | 0 | 0 | 0 | 0 | | 0 | 2 | 1 | 0 | 0 | 0 | 0 | 33 | 0 | 0 | 10 | 0 | 26 | 4 | 0 |
| 20 < %GC < 30 | | 1 | 0 | 0 | 0 | 2 | | 2 | 1 | 5 | 1 | 0 | 0 | 0 | 5 | 1 | 2 | 1 | 0 | 0 | 42 | 0 |
| GC% < 20 | | 3 | 0 | 3 | 0 | 0 | | 6 | 0 | 2 | 1 | 0 | 0 | 0 | 3 | 1 | 24 | 5 | 0 | 1 | 5 | 0 |

**Figure S12: GC content of complete copies of the main subfamilies of Transposable Elements (TEs) in the genomes of the *B. cinerea* strains B05.10, Sl3 and Vv3.** Each symbol in the graph corresponds to one copy of a TE or to several copies with the same GC content. The table indicates the total number of complete copies in each genome, as well as the number of copies in different ranges of GC content.

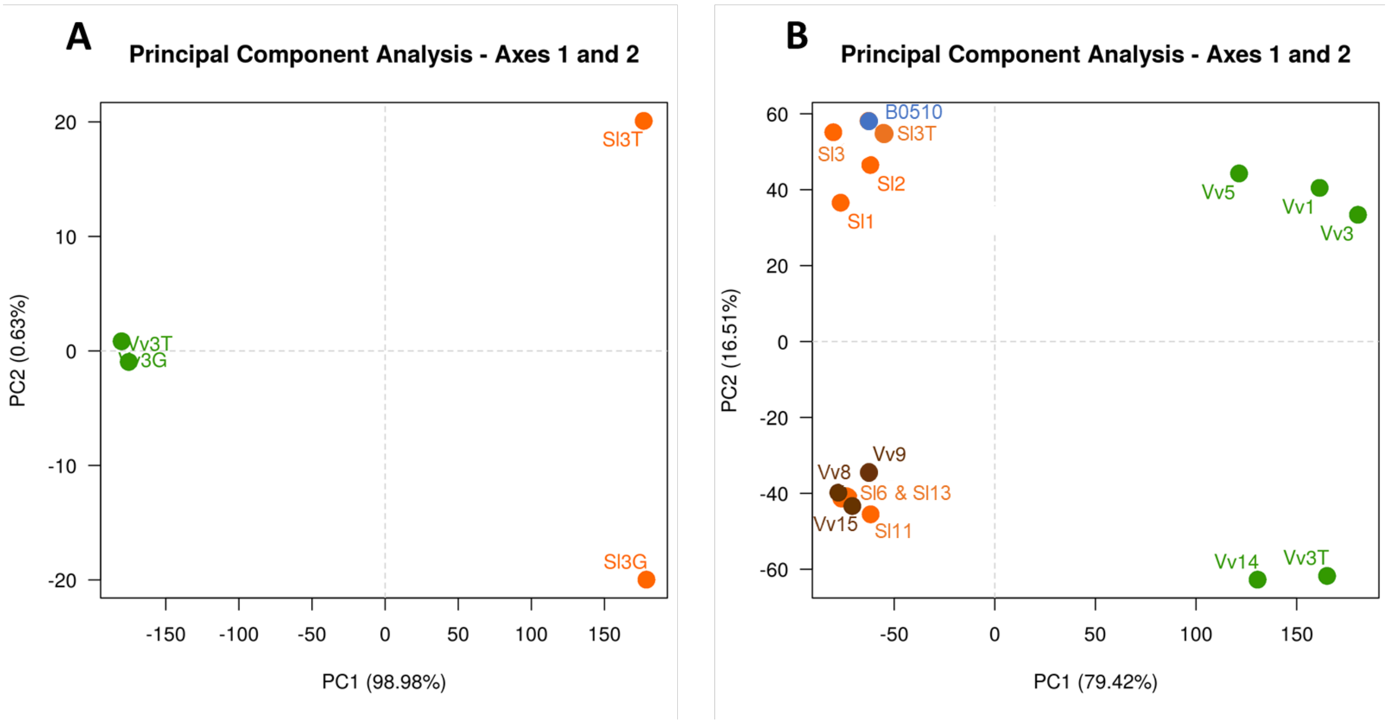


**Figure S13: Principal Component Analysis (PCA) of the repertoires of small RNAs in different strains of *B. cinerea*. A.** Vv3 and Sl3 strains were cultivated either on grape juice (Vv3G and Sl3G), either on tomato juice (Vv3T and Sl3T) medium. **B**. The analysis was extended to five additional strains from the T population (in orange), three additional strains from the G1 population (in green), three strains from the G2 population (in brown), and the B05.10 strain, all grown on grape juice medium.

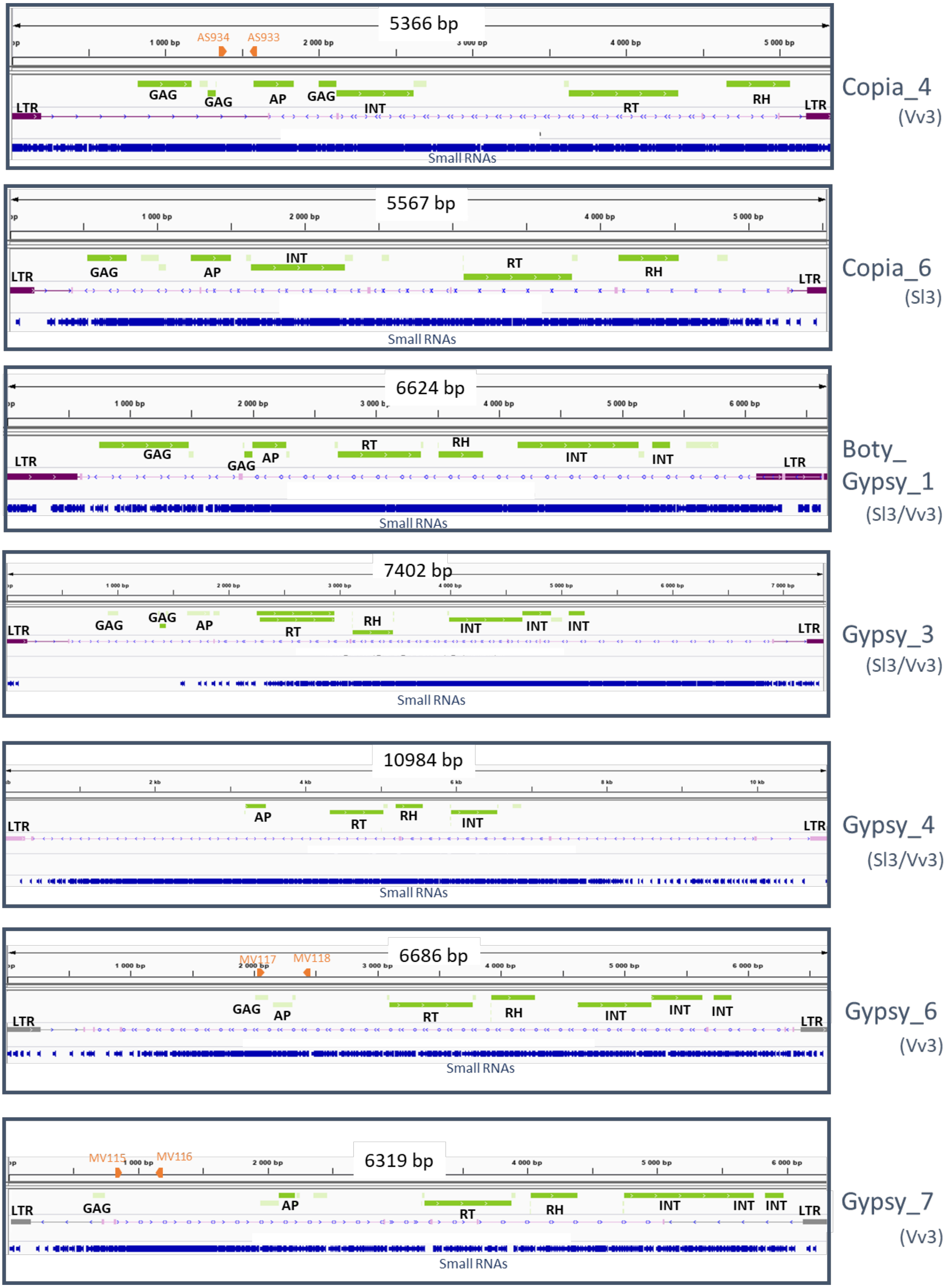

5366 bp
AS934
AS933
GAG
GAG
AP
GAG
INT
RT
RH
LTR
LTR
Small RNAs
Copia_4
(Vv3)
5567 bp
LTR
GAG
AP
INT
RT
RH
LTR
Small RNAs
Copia_6
(Sl3)
6624 bp
LTR
GAG
GAG
AP
RT
RH
INT
INT
LTR
Small RNAs
Boty_
Gypsy_1
(Sl3/Vv3)
7402 bp
LTR
GAG
GAG
AP
RT
RH
INT
INT
INT
LTR
Small RNAs
Gypsy_3
(Sl3/Vv3)
10984 bp
LTR
AP
RT
RH
INT
LTR
Small RNAs
Gypsy_4
(Sl3/Vv3)
6686 bp
MV117
MV118
LTR
GAG
AP
RT
RH
INT
INT
INT
LTR
Small RNAs
Gypsy_6
(Vv3)
6319 bp
MV115
MV116
LTR
GAG
AP
RT
RH
INT
INT
INT
LTR
Small RNAs
Gypsy_7
(Vv3)

**Figure S14: Retrotransposons that generate_small RNAs in the *B. cinerea* strains Sl3 and/or Vv3.** Transposable Elements (TEs) were annotated using the REPET package (http://urgi.versailles.inra.fr/tools/REPET; Amselem et al., 2015) and classified according to their structure and sequence similarities to characterized eukaryotic TEs (Wicker et al., 2007, Hoede et al., 2014). LTR: Long terminal repeats. GAG: Capsid protein. AP: Aspartic proteinase. RT: Reverse transcriptase. RH: RNAse H. INT: Integrase. Small RNAs were mapped to the TE consensus and displayed in blue. The primers used in this study are indicated in orange.

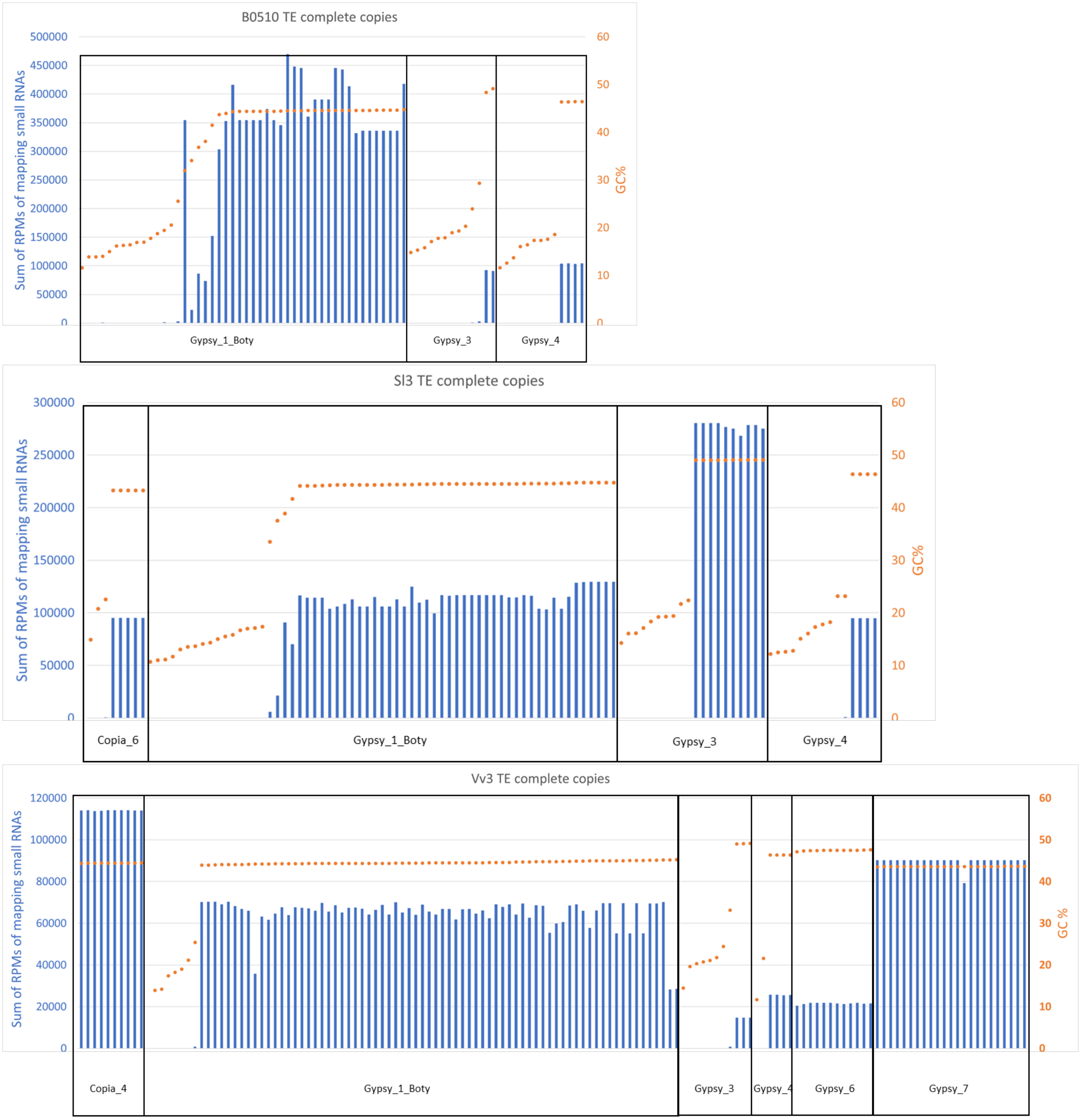


**Figure S15: Mapping of small RNA reads on complete copies of TEs in relation to their GC percent**. For each complete copy of TE, the total RPM of mapping small RNAs reads is represented as a blue bar whereas the GC percent is indicated as an orange dot. Only complete copies with high GC content produce significant amounts of small RNAs.

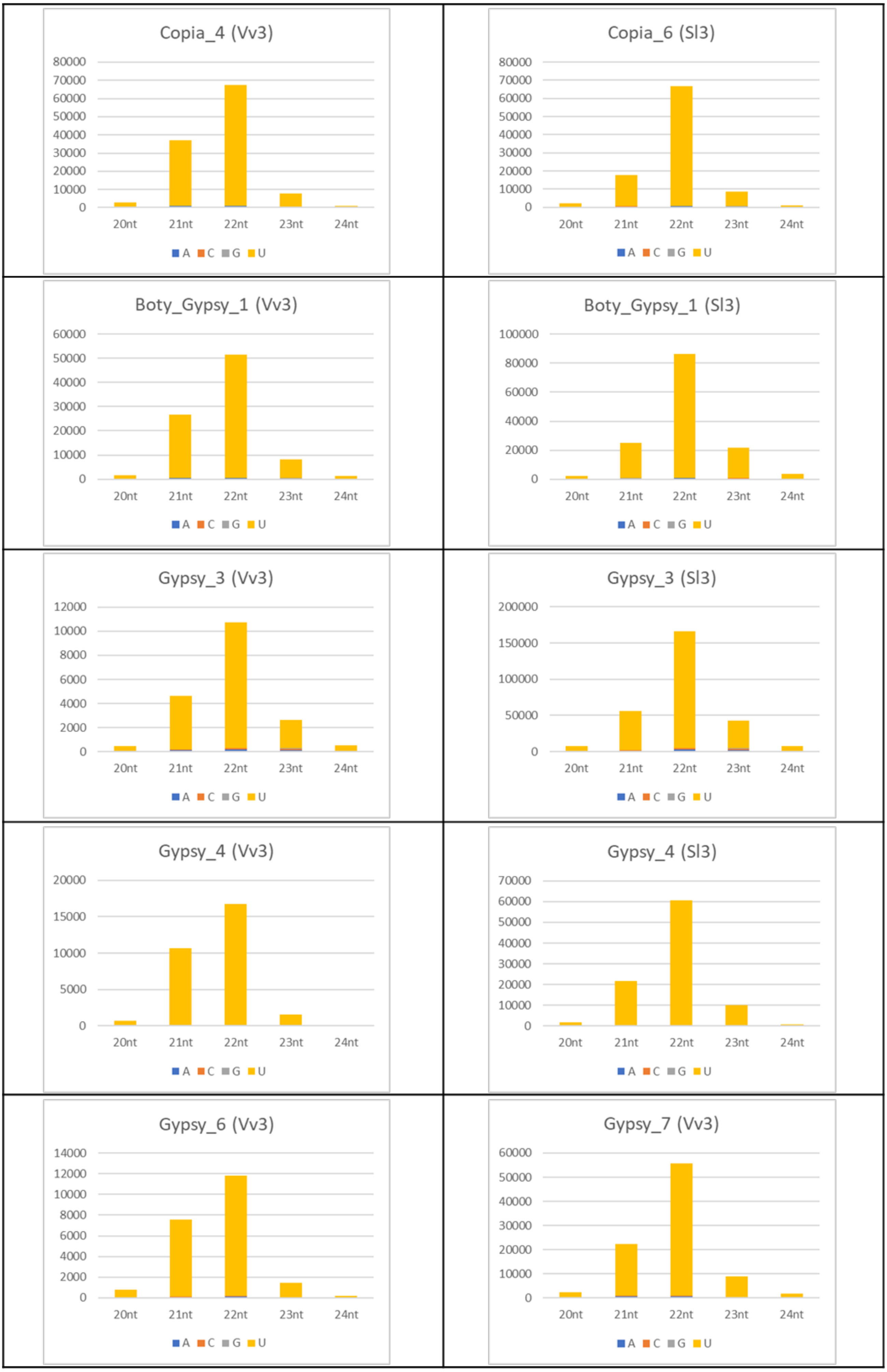
Copia_4 (Vv3)
80000
70000
60000
50000
40000
30000
20000
10000
0
20nt
21nt
22nt
23nt
24nt
A C G U
Copia_6 (Sl3)
80000
70000
60000
50000
40000
30000
20000
10000
0
20nt
21nt
22nt
23nt
24nt
A C G U
Boty_Gypsy_1 (Vv3)
60000
50000
40000
30000
20000
10000
0
20nt
21nt
22nt
23nt
24nt
A C G U
Boty_Gypsy_1 (Sl3)
100000
80000
60000
40000
20000
0
20nt
21nt
22nt
23nt
24nt
A C G U
Gypsy_3 (Vv3)
12000
10000
8000
6000
4000
2000
0
20nt
21nt
22nt
23nt
24nt
A C G U
Gypsy_3 (Sl3)
200000
150000
100000
50000
0
20nt
21nt
22nt
23nt
24nt
A C G U
Gypsy_4 (Vv3)
20000
15000
10000
5000
0
20nt
21nt
22nt
23nt
24nt
A C G U
Gypsy_4 (Sl3)
70000
60000
50000
40000
30000
20000
10000
0
20nt
21nt
22nt
23nt
24nt
A C G U
Gypsy_6 (Vv3)
14000
12000
10000
8000
6000
4000
2000
0
20nt
21nt
22nt
23nt
24nt
A C G U
Gypsy_7 (Vv3)
60000
50000
40000
30000
20000
10000
0
20nt
21nt
22nt
23nt
24nt
A C G U

**Figure S16: Small RNAs produced by the *B. cinerea* strains Sl3 and Vv3: Size distribution and 5' nucleotide.** Counting of small RNAs mapping each of the seven small RNAs-producing retrotransposons are indicated by Reads Per Million (RPMs). Only the reads of length 20-24nt with at least five RPMs in at least one of the Vv3 or Sl3 library were considered here.

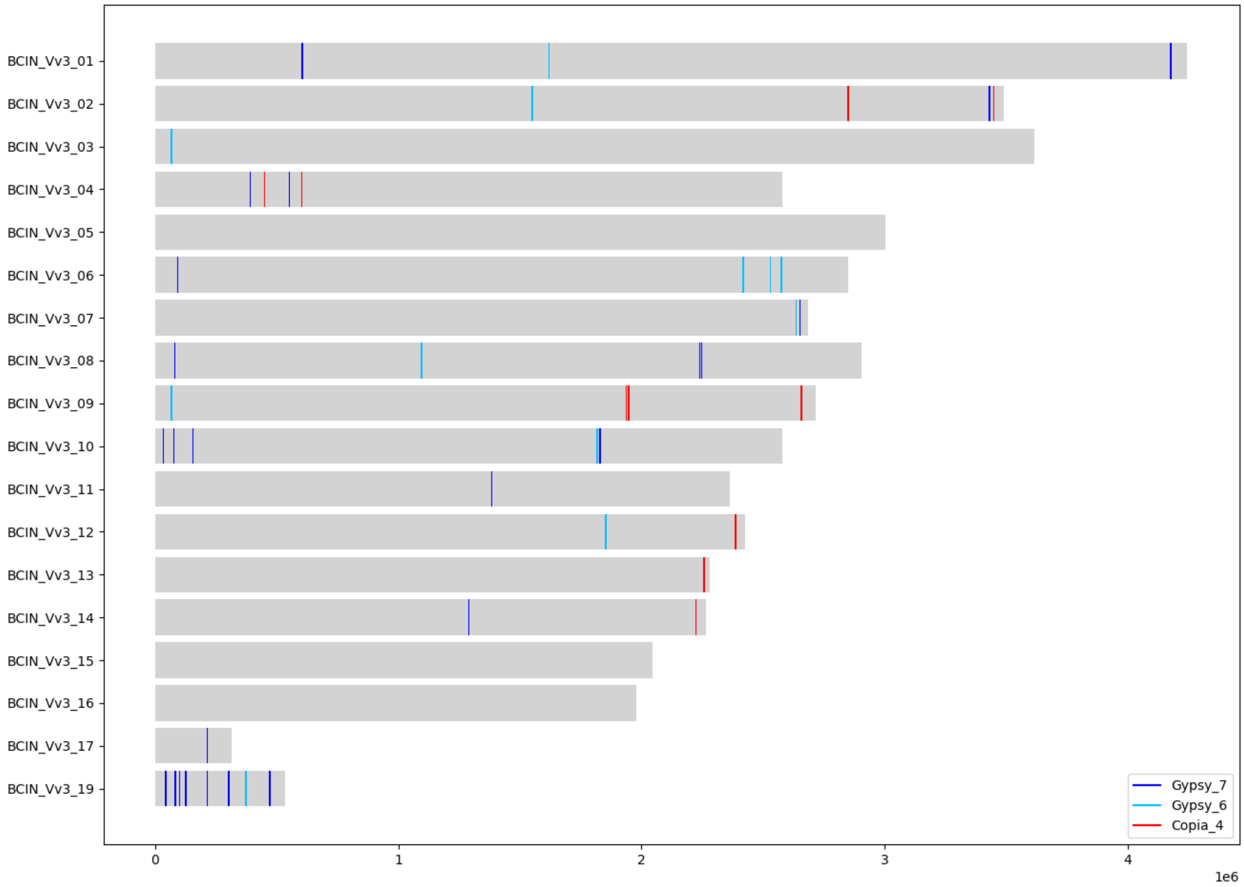


**Figure S17: Positions of complete copies of the TE Copia_4, Gypsy_6 and Gypsy_7 in the *B. cinerea* Vv3 genome.**

Copia_4

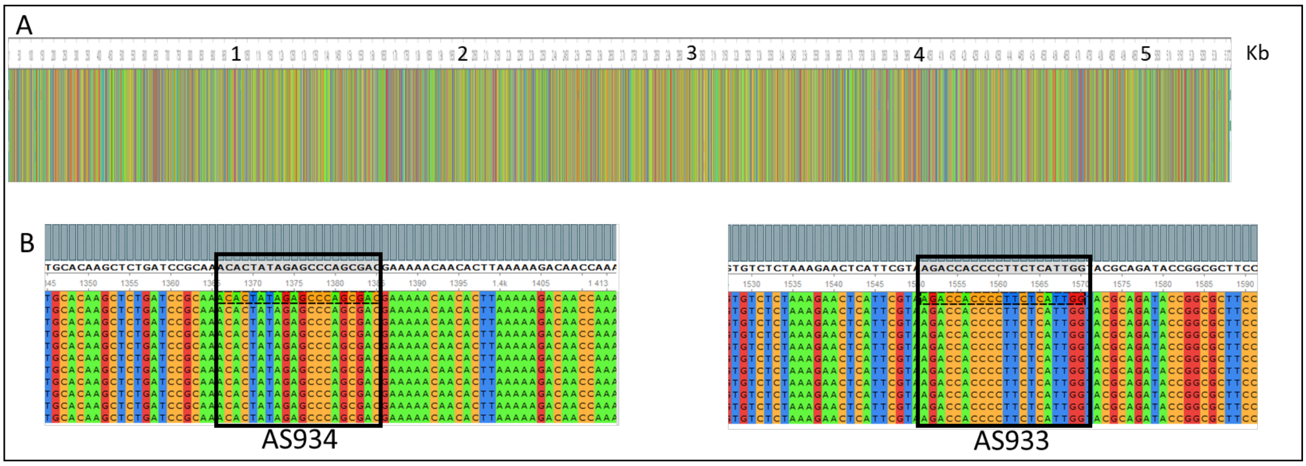


Gypsy_6

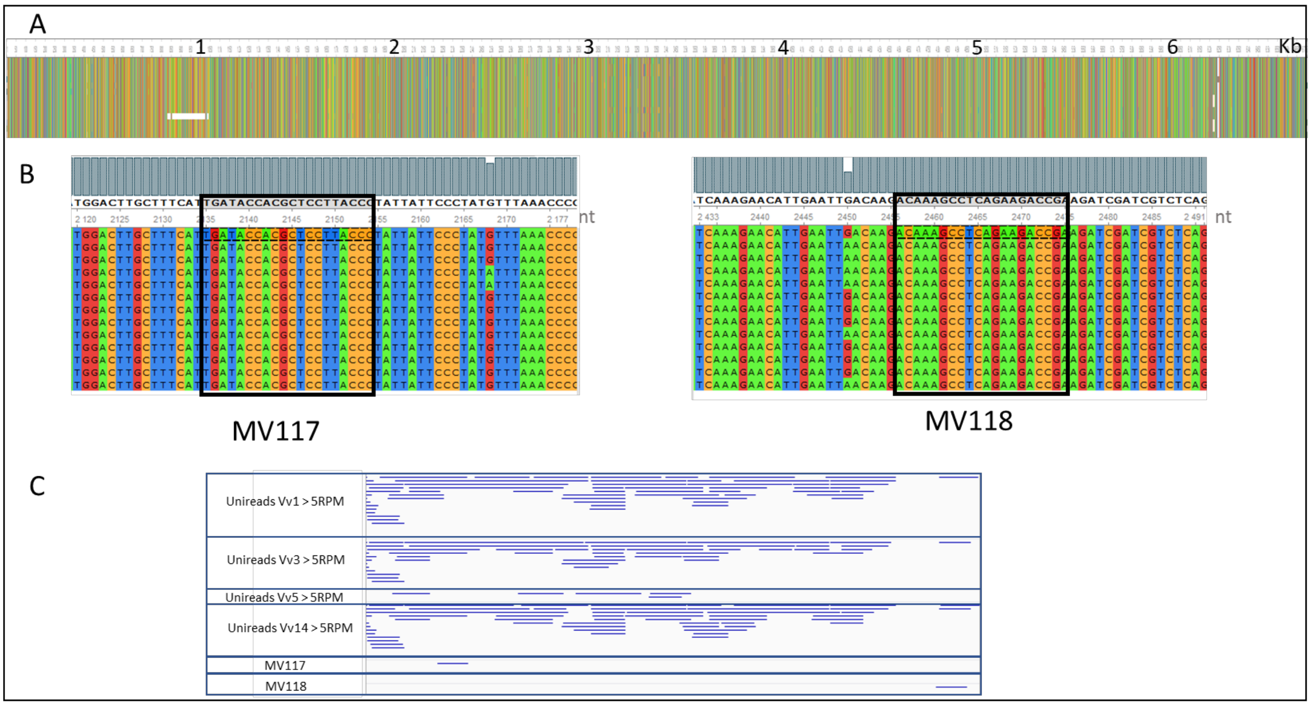


Gypsy_7

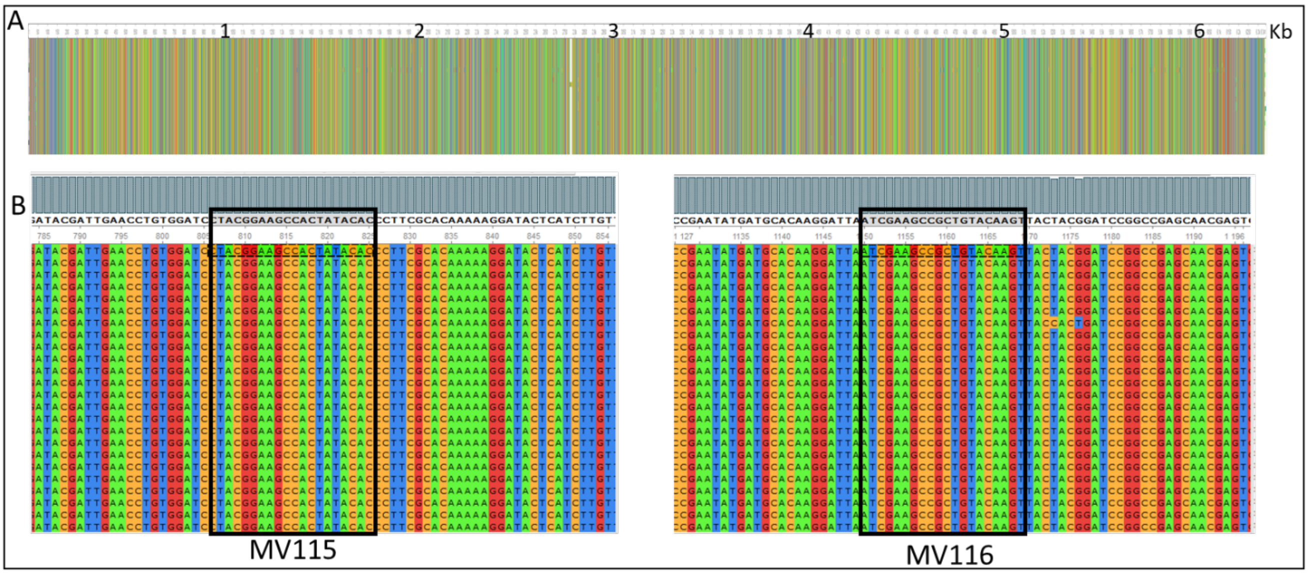

**Figure S18: Alignment of Copia_4, Gypsy_6 and Gypsy_7 consensus sequences with their respective complete copies in the Vv3 genome. A.** Alignment of the whole sequences. **B.** Zoom on the regions where the PCR primers were designed. **C.** Mapping of Vv1, Vv3, Vv5 and Vv14 small RNA unireads (>5RPM) on Gypsy_6 consensus and position of the primers (MV117, MV118) used for PCR.
